# Quantification of spatial and phenotypic heterogeneity in an agent-based model of tumour-macrophage interactions

**DOI:** 10.1101/2022.05.26.493564

**Authors:** Joshua A. Bull, Helen M. Byrne

## Abstract

We introduce a new spatial statistic, the weighted pair correlation function (wPCF). The wPCF extends the existing pair correlation function (PCF) to describe spatial relationships between points marked with combinations of discrete and continuous labels. We validate its use through application to an agent-based model (ABM) which simulates interactions between macrophages and tumour cells. These interactions are influenced by the spatial positions of the cells and by macrophage phenotype, a continuous variable that ranges from anti-tumour to pro-tumour. By varying model parameters that regulate macrophage phenotype, we show that the ABM exhibits behaviours which resemble the ‘three Es of cancer immunoediting’: Equilibrium, Escape, and Elimination.

We use the wPCF to analyse synthetic images generated by the ABM. We show that the wPCF generates a ‘human readable’ statistical summary of where macrophages with different phenotypes are located relative to both blood vessels and tumour cells. In combination with the cross-PCF (describing interactions between vessels and tumour cells), we show further that each of the three Es of immunoediting is characterised by a distinct ‘PCF signature’. By applying dimension reduction techniques to this signature, we identify its key features and train a support vector machine classifier to distinguish between simulation outputs based on their PCF signature. This proof-of-concept study shows how multiple spatial statistics can be combined to analyse the complex spatial features that the ABM generates, and to partition them into interpretable groups.

The intricate spatial features produced by the ABM are similar to those generated by state-of-the-art multiplex imaging techniques which distinguish the spatial distribution and intensity of multiple biomarkers in biological tissue regions. Applying methods such as the wPCF to multiplex imaging data would exploit the continuous variation in biomarker intensities and generate more detailed characterisation of the spatial and phenotypic heterogeneity in tissue samples.

**Author summary:** Multiplex images provide exquisitely detailed information about the spatial distribution and intensity of up to 40 biomarkers within two-dimensional tissue regions, creating challenges and opportunities for quantitative analysis. Although stain intensities are measured on a continuous scale, they are typically converted into discrete labels to simplify subsequent spatial analysis. In this paper we propose a new spatial statistic, the weighted pair correlation function (wPCF), which exploits, rather than neglects, the continuous variation in stain intensity contained in multiplex images, and can characterise both spatial and phenotypic heterogeneity.

As proof-of-principle, we apply the wPCF to synthetic data that resemble multiplex images of solid tumours. We generate data from an agent-based model (ABM) that simulates macrophage-tumour interactions. The wPCF shows how the continuous label describing macrophage phenotype is spatially related to categorical labels associated with tumour cells and blood vessels. We demonstrate that correlation functions can categorise spatial relationships in a manner which is interpretable and quantitative.

The methods we present can be used to analyse both ABM simulations and multiplex imaging data, with applications that go beyond macrophage phenotype to include other biological processes that exhibit continuous variation (e.g., cancer cell stemness, biomarkers for T-cell exhaustion, and levels of oxygenation).

## Introduction

Tumours are highly heterogeneous structures, containing diverse populations of tumour cells, blood vessels, stromal cells and immune cells. The immune landscape within solid tumours is complex and varied [1, 2], with both innate and adaptive immune cells implicated in pro- and anti-tumour responses [3]. For example, high densities of tumour associated macrophages have been associated with poor prognosis in breast, prostate and head and neck cancer and with good prognosis in colorectal and gastric cancer [4, 5]. These differences may be due to the relative numbers of pro-tumour (‘M_1_’) and anti-tumour (‘M_2_’) macrophages in these cancers, but they may also be due to their morphology and spatial distribution [6–9]. For example, in non-small cell lung cancer, high infiltration rates of M_1_ macrophages into tumour islets, but not tumour stroma, have been associated with increased patient survival [10].

While macrophages are often classified as either M_1_ or M_2_, individual macrophages may exhibit a variety of behaviours. Further, their overall behaviour, or phenotype, may change over time in response to multiple microenvironmental cues [11]. Macrophage phenotype is often defined in terms of expression levels of multiple functional markers such as CD68, CD163, CD204 and CD206 [9]. It is difficult to resolve this level of heterogeneity using traditional immunohistochemistry (IHC), which typically permits only one or two markers per image. By contrast, multiplex imaging modalities, such as multiplexed IHC and imaging mass cytometry (IMC), can map expression levels of up to 40 different cellular markers and, as such, resolve the spatial position and phenotype of individual cells, including macrophages [7, 12–14].

In Fig 1A we present a typical multiplex image which shows spatial variation in the intensity levels of three macrophage markers (CD68, CD163 and CD206), reproduced from [12]. In Fig 1B we show how the average intensity levels of these markers are used to classify segmented cells. Defined threshold intensities are used to determine whether cells are negative (−), positive (+) or extremely positive (++) for a particular marker. The classifications are combined to identify seven macrophage subtypes: CD68+, CD68++CD163+, CD68+CD163+, CD68+CD163+CD206+, CD68+CD206+, CD68+CD206++, and CD68+IRF8+. By associating one of these classifications with the centroid of each macrophage, a spatial map of macrophage subtypes can be generated and used for subsequent spatial analyses. Such analyses may include correlations of cell densities, measurements of distances between cells, or more complex spatial statistics such as pair correlation functions (PCFs) which account for spatial relations between points [15]. Fig 1C shows how the average intensity levels of CD68, CD163 and CD206 associated with each macrophage subtype vary across *n* = 35 patients (data reproduced from [12]). Variation in marker levels occurs both within a tissue sample and across patients.

**Fig 1.**
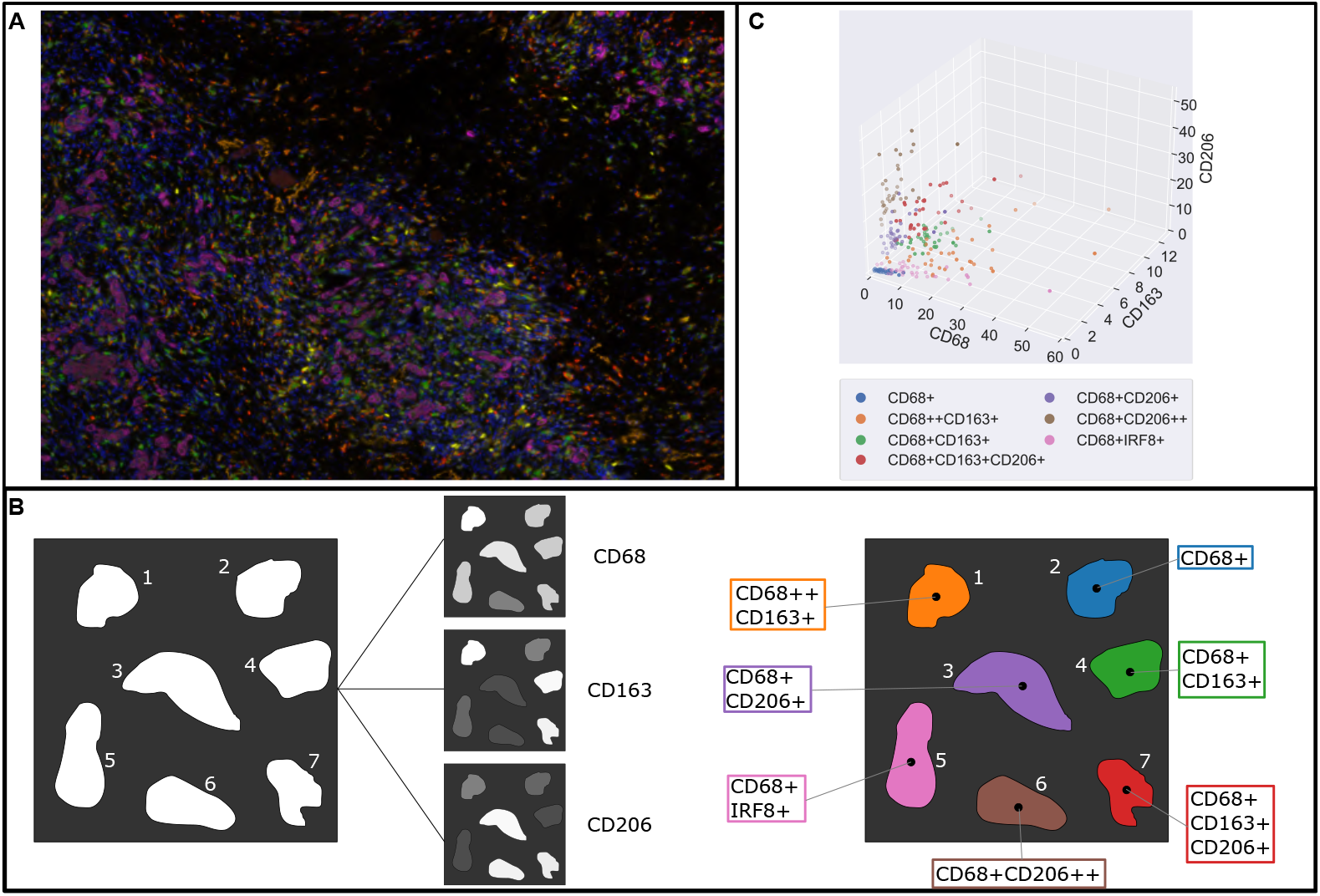
Typical process for analysing macrophage phenotypes in multiplex imaging. A: A multiplex image showing macrophages of varying phenotypes, reproduced from Fig 1c in [12]. Expression levels of different functional markers (e.g., CD68, CD163, CD206) are shown as differing intensities of separate stains, visualised as a false-colour image. B: Schematic indicating how continuous stain intensities represented in a multiplex image are converted into different categories indicating macrophage phenotype. In this example, cell colours in the multiplex image are converted into stain intensities for CD68, CD163 and CD206. Thresholds are assigned to each stain to distinguish whether a cell is negative, positive, or extremely positive, for each stain. Each macrophage is then assigned one of 7 different potential phenotypes, based on combinations of positivity or negativity for each stain. C: Data reproduced from Fig 1h in [12]. Points represent the average stain intensity of CD68, CD163, and CD206 measured in macrophages assigned to each of the seven phenotype subtypes in each of n=35 patients. Note that macrophages from the same subtype in different patients have differing levels of intensity of each marker, so the same categorical label may be applied to macrophages with a wide range of continuous expression levels.

Stratification of macrophage populations into discrete classes (e.g., M_1_ and M_2_ classes, or the seven subtypes shown in Fig 1), neglects the full range of information available from multiplex images. In this paper, we show how resolving continuous variation in cell labels clarifies the relationship between macrophage phenotype and spatial heterogeneity in solid tumours. To achieve this, we introduce the weighted pair correlation function (wPCF), a new spatial statistic which accounts for continuous variation in labels such as cell subtype, phenotype, or marker expression levels. While similar ‘marked point patterns’ have been studied in ecology [16–18] and astronomy [19, 20], few methods consider the spatial correlation of continuous marks. Existing methods, such as Stoyan’s mark correlation function [16, 19, 21, 22] or the mark variogram [18, 23], generally depend only on the distance *r* between point pairs and attempt to quantify the correlation between marks at distance *r*. In biological applications, a more pertinent problem is to determine whether points with a particular mark, or range of marks, are correlated at distance *r*. The wPCF addresses this, identifying spatial interactions between points of one type and those whose mark matches one of a range of target values.

We test and validate the wPCF using synthetic data generated from a two-dimensional agent-based model (ABM) of tumour growth that accounts for tumour-macrophage interactions and dynamic changes in macrophage phenotype. ABMs are well-suited for generating labelled point pattern data, since each cell is represented by an agent whose behaviour is determined by subcellular variables (describing, for example, cell cycle state [24] or phenotype), and its interaction with the environment (e.g., through force laws describing mechanical interactions between cells). These subcellular variables can be used to represent marker expression levels, meaning that each agent is associated with a point representing its cell centre together with a collection of continuous or categorical labels. Data from such ABMs can be analysed using PCFs [25–30] or cross-PCFs [31, 32], an extension of the PCF which accounts for interactions between cells of different types.

The off-lattice, force-based ABM that we develop is motivated by an experimental study by Arwert *et al* [33] which investigates how macrophage phenotype depends on spatial location relative to a tumour mass and nearby vasculature, and how the spatial distribution of the different macrophage phenotypes influences the tumour’s growth dynamics. In brief, anti-tumour macrophages extravasate from blood vessls and migrate towards clusters of tumour cells, in response to tumour-derived signals such as colony stimulating factor-1 (CSF-1). Exposure to TGF-*β* in the tumour increases macrophage sensitivity to C-X-C chemokine ligand type 12 (CXCL12) and drives them towards a pro-tumour phenotype. At the same time, CXCL12 produced by perivascular fibroblasts biases the movement of these M_2_ macrophages towards neighbouring blood vessels. As they migrate out of the tumour, the pro-tumour macrophages express epidermal growth factor (EGF), a tumour cell chemoattractant. In this way, M_2_ macrophages facilitate metastasis by guiding the tumour cells towards the vasculature [33, 34].

The hybrid ABM developed in this paper builds on existing differential equation models [35, 36] and ABMs [36, 37] that focus on specific tumour-macrophage interactions, such as the CSF-1/EGF paracrine loop that mediates cross-talk between tumour cells and macrophages. Our model accounts for macrophage extravasation in response to tumour-derived CSF-1, their subsequent tumour infiltration, and the CSF-1/EGF paracrine loop that mediates cross-talk between tumour cells and macrophages. Models of macrophage-tumour interactions often view macrophages as a homogeneous population [38, 39], or account for multiple macrophage subtypes (typically M_1_ and M_2_) [40–43] and their interactions with T-cells [44–46]. Eftimie [47] and El-Kenawi *et al*. [48] have developed models that view macrophage phenotype as a continuous variable whose dynamics are governed by environmental cues, such as pH. We represent macrophage phenotype as a continuous variable, *p*, whose dynamics depend on local levels of TGF-*β* and determine macrophage behaviour.

We vary ABM parameters, and show that it generates a range of spatial patterns and qualitative behaviours that resemble ‘The Three Es of cancer immunoediting’ [49]. In particular, as we vary model parameters associated with the rate of macrophage extravasation and their chemotactic sensitivity to CSF-1, we observe tumour Elimination, Escape, and Equilibrium. We analyse simulation outputs at a fixed timepoint using wPCFs and cross-PCFs. We explain how, taken together, wPCFs and cross-PCFs provide a description of simulation outputs which is both quantitative and interpretable. We then show how the spatial statistics can be combined and analysed using principal component analysis (PCA) to identify the key features that characterise Elimination, Escape, and Equilibrium for our ABM simulations. We show further how the principal components can be used to classify unseen data from ABM simulations. More generally, this study serves as a powerful proof of concept: it shows how combinations of wPCFs and cross-PCFs could be used to accurately classify the complex spatial and phenotypic patterns formed by cells in multiplex images, without manual thresholding of marker intensities.

The remainder of the paper is structured as follows. We first describe our ABM, emphasising those features which make it appropriate for generating synthetic imaging data. We introduce the wPCF and illustrate how it can be interpreted as a series of cross-PCFs which vary continuously with the label of interest, here macrophage phenotype. We then apply the wPCF to synthetic data generated from our ABM, and show that it provides a more detailed description of the relationship between macrophage phenotype and spatial location than cross-PCFs. We define a ‘PCF signature’, consisting of two wPCFs and a cross-PCF, which describes the spatial relationships between blood vessels, tumour cells, and macrophages of each phenotype. The signature can be interpreted as a high-dimensional vector, and we apply principal component analysis (PCA) to reduce its dimensionality. We demonstrate a proof-of-concept classification algorithm by using the first 100 principal components to train a simple classifier which distinguishes between simulation outputs that correspond to the three Es of immunoediting (i.e., Escape, Equilibrium and Elimination). Finally, we calculate PCF signatures for dynamic ABM data, and show that a single simulation may transition between tumour Equilibrium, Escape and Elimination at different timepoints.

## Materials and Methods

### Agent-based model (ABM)

We present a 2D, multiscale, off-lattice ABM which extends an existing model of macrophage infiltration into tumour spheroids by accounting for continuous and dynamic variation in macrophage phenotype [30, 50]. The new model simulates a growing tumour embedded in a small tissue region *in vivo*, and includes phenotype-dependent interactions between macrophages, blood vessels and tumour cells. We outline the ABM here, and refer the interested reader to S1 Appendix: Model Description for details of the implementation and default parameter values. The ABM is implemented within the open source Chaste (Cancer, Heart and Soft Tissue Environment) modelling environment [51–53].

### Overview

The ABM distinguishes four cell types: stromal cells, tumour cells, necrotic cells, and macrophages. Their dynamics are influenced by five diffusible species: oxygen (*ω*), CSF-1 (*c*), CXCL12 (*ξ*), TGF-*β* (*g*) and EGF (*ϵ*). In the 2D Cartesian geometry, blood vessels are represented by fixed points which do not compete for space with the cell populations and which act as distributed sources of oxygen [54]. A schematic of the ABM is presented in Fig 2.

**Fig 2.**
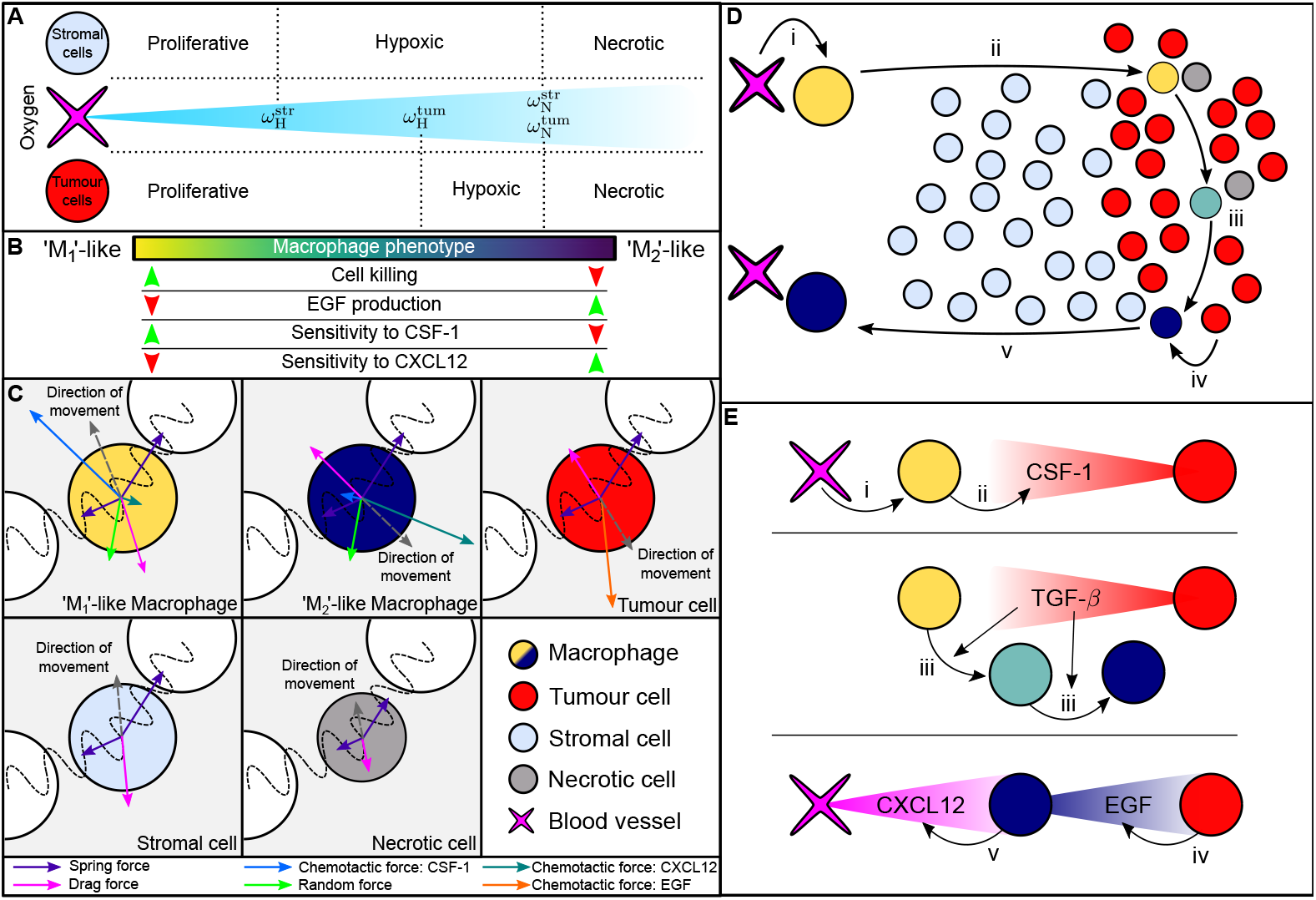
Schematic summarising the key interactions that are included in the agent based model. A: Oxygen is supplied by blood vessels and consumed by stromal cells and tumour cells. Cell-cycle progression is determined by a cell’s local oxygen concentration: a cell may be ‘proliferative’ (and progress through its cell cycle), ‘hypoxic’ (the cell cycle is temporarily paused until oxygen concentrations return to a sufficiently high level) or ‘necrotic’ (the cell becomes necrotic cell and degrades). Cell cycles also pause if there is insufficient space available for proliferation. B: Macrophage behaviour depends on phenotype *p*, modulating their rates of tumour cell killing, EGF production, and chemotactic sensitivity to gradients of CSF-1 and CXCL12. C: Forces acting on different cell types. Macrophages are subject to mechanical forces due to interactions with nearby cells, and random forces which simulate their exploration of their environment as highly motile cells. Macrophages also experience chemotactic forces that are directed up spatial gradients of CSF-1 and CXCL12, and whose magnitude depends on *p*. Tumour cells experience mechanical forces due to interactions with neighbouring cells, and chemotactic forces in the direction of increasing EGF. Stromal cells experience mechanical forces due to interactions with neighbouring cells. Necrotic cells experience these interaction forces, which decrease in magnitude as they decrease in size. All cells experience a drag force. D: Summary of the phases of macrophage-mediated tumour cell migration in our ABM. i) M_1_ macrophages extravasate from blood vessels in response to CSF-1. ii) M_1_ macrophages migrate into the tumour mass in response to CSF-1, where they may kill tumour cells. iii) Exposure to TGF-*β* causes macrophages to adopt an M_2_ phenotype. iv) M_2_ macrophages produce EGF, which acts as a chemoattractant for tumour cells. v) M_2_ macrophages migrate towards blood vessels, in response to CXCL12 gradients. E: Schematic summarising the sources of CSF-1, TGF-*β*, EGF and CXCL12 in our model, and their interactions with cells, as described in steps i-v of panel D.

Following [55], critical oxygen thresholds for hypoxia (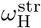 and 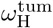) and necrosis (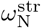 and 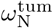) relate the rates of cell cycle progression of stromal and tumour cells to local oxygen levels (see Fig 2A for details). For example, if, at time *t* > 0, 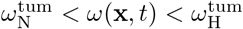, then the cell cycle of a tumour cell at position **x** will immediately halt and remain paused until the local oxygen concentration rises above the tumour hypoxic threshold. If, however, the oxygen concentration falls below the tumour necrosis threshold, then the cell becomes necrotic (this switch is irreversible). Necrotic cells occupy space for a finite time period during which their size decreases linearly to zero and they are then removed from the simulation. Blood vessels also act as entry points for macrophages, which infiltrate the tissue and alter their phenotype (and, hence, behaviour) at rates which depend on local levels of TGF-*β* (see Fig 2B).

We represent each cell by the spatial coordinates of its centre of mass and determine its movement by balancing the forces that act on each it. Using an overdamped form of Newton’s second law and neglecting inertial terms, we have that for cell *i*

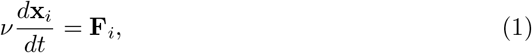

where *ν* is the assumed constant drag coefficient and **F**_*i*_ denotes the net force acting on cell *i* at position **x**_*i*_ and time *t*. The forces that act on a cell depend on its type (see Fig 2C and S1 Appendix: Model Description). Cells interact via spring forces if their centres are within a distance *R*_int_ of each other [56]; intercellular adhesion and volume exclusion are represented by attractive and repulsive forces respectively. We also associate with each cell an approximate area, and stromal cells which are so mechanically compressed that their area falls below a threshold proportion 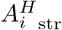 of their normal area pause their cell cycle due to contact-inhibition (see S1 Appendix: Model Description for details).

### Macrophage phenotype

Since [33] describes a unidirectonal transition of macrophage phenotype, we use a single continuous subcellular variable to represent macrophage phenotype. This variable, *p* ∈ [0, 1], determines macrophage behaviour, with *p* < 0.5 denoting a primarily anti-tumour M_1_ phenotype and *p* > 0.5 representing a primarily pro-tumour M_2_ phenotype. While M_1_ and M_2_ describe two broad categories of macrophage, their underlying behaviour is dependent on the value of *p* rather than this categorisation, and for simplicity we may refer to macrophages whose phenotype is close to 0.5 as ‘transitioning’ macrophages. We assume that, following extravasation, macrophage *i* has a phenotype *p*_*i*_ = 0. Macrophage exposure to TGF-*β* levels above a threshold value, *g*_crit_, causes *p*_*i*_ to increase at a constant rate Δ*p*, per timestep *dt*, until the maximum value *p*_*i*_ = 1 is reached and the macrophage has a fully M_2_ phenotype. Its phenotype remains fixed at *p*_*i*_ = 1 for all later times. Thus, we have:

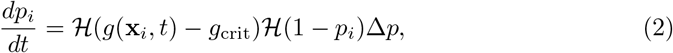

where 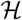 is the Heaviside function (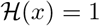 when *x* > 0 and 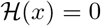 otherwise).

We now explain how changes in phenotype *p* affect macrophage behaviour and function, and how these changes are incorporated into the ABM (see also Fig 2B).

#### Macrophage extravasation

Macrophages enter the domain via blood vessels with a probability per hour *P*_ex_ which is an increasing, saturating function of CSF-1:

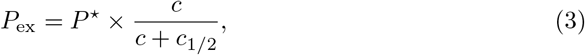

where the non-negative parameter *P*^⋆^ represents the maximum probability per hour of macrophage extravasation from a vessel, and *c*_1/2_ is the concentration of CSF-1 at which the probability is half-maximal.

#### Macrophage chemotactic forces

Fig 2C shows the forces which act on different cell types (functional forms for these forces are given in S1 Appendix: Model Description). Here we highlight two macrophage-specific forces which describe their directed movement up spatial gradients of CSF-1 and CXCL12, and whose magnitude varies with phenotype *p*. Noting that M_1_ macrophages are sensitive to CSF-1 and insensitive to CXCL12 (and conversely for M_2_ macrophages), we assume that chemotactic forces depend linearly on phenotype. The chemotactic forces acting on macrophage *i*, at position **x**_*i*_ with phenotype *p*_*i*_, are therefore:

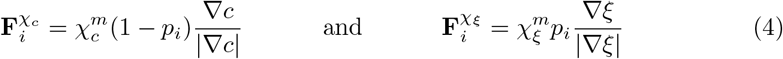

respectively, where the non-negative parameters 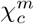 and 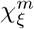 indicate macrophage sensitivity to spatial gradients of CSF-1 and CXCL12, and ∇*c* and ∇*ξ* are evaluated at **x**_*i*_. The forces 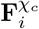 and 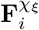 contribute to the net force **F**_*i*_ in Eq (1) (see Fig 2C and S1 Appendix: Model Description).

#### Macrophage cell killing

We assume that when a macrophage and a tumour cell are within the interaction radius *R*_int_ then the macrophage will attempt to kill the tumour cell, with M_1_ macrophages more likely to kill a tumour cell than M_2_ macrophages. We define a probability of cell kill per hour, *P*_*φ*_, which is a monotonic decreasing function of *p*. We suppose further that, after a macrophage has killed a tumour cell, it experiences a ‘cooldown’ period of *t*_cool_ hours during which it cannot attempt tumour cell killing. Thus, we associate with macrophage *i* a subcellular timer *t*_*φ,i*_ that is updated in real time and set to zero on tumour cell killing. We define *P*_*φ,i*_ as:

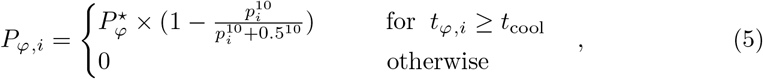

where 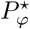 is the maximum probability of tumour cell killing. If macrophage *i* is sufficiently close to attack multiple tumour cells then one is selected at random for cell death. Killed tumour cells are labelled as ‘necrotic’ and decay in the same way as other necrotic cells.

#### Macrophage production of EGF

The diffusible cytokine EGF, *ϵ*, is produced by M_2_ macrophages and undergoes natural decay. It is also a potent chemoattractant for tumour cells. For simplicity, we assume that macrophage *i* produces EGF at a rate which is linearly proportional to its phenotype *p*_*i*_, with constant of proportionality *κ*_*ϵ*_. Denoting by *D*_*ϵ*_ and *λ*_*ϵ*_ the assumed constant diffusion coefficient and natural decay rate of EGF, we suppose that its evolution can be described by the following reaction diffusion equation:

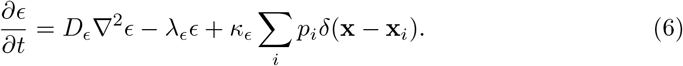

where *δ*(**x**) = **1** when **x** = 0 and *δ*(**x**) = **0** elsewhere. In (6), we sum over all macrophages to determine the net rate of production at spatial position **x**.

### Spatial statistics

In order to compute spatial statistics, we introduce the following notation. Consider an object *i* (which may be a cell or a blood vessel), whose centre is located at **x**_*i*_ = (*x*_*i*_, *y*_*i*_) at time *t*. We associate with object *i* a categorical label *q*_*i*_ ∈ {*B, M, S, T, N*} which indicates whether it is a blood vessel, macrophage, stromal cell, tumour cell or necrotic cell. Given a target label *Q* ∈ {*B, M, S, T, N*}, the binary target function Θ(*Q, q*_*i*_) indicates whether the label *q*_*i*_ matches *Q*:

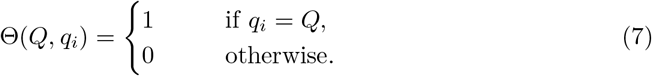

### Cross pair correlation function (cross-PCF)

The cross-PCF identifies spatial correlations between objects with categorical labels that are separated by a distance *r*. We define a sequence of annuli, of inner radius *r*_*k*_ and outer radius *r*_*k*_ + *dr* where *r*_0_ = 0 and *dr* > 0. We denote by 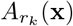 the area of the annulus with inner radius *r*_*k*_ that is centred at the point **x**. If this annulus lies wholly inside the domain then 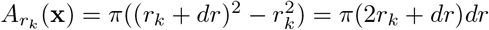; otherwise, only the area contained within the domain is recorded. The indicator function, *I*_*k*_(*r*), is

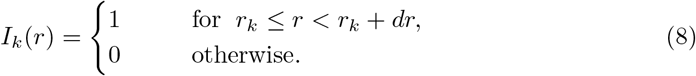

We calculate the cross-PCF for blood vessels and tumour cells by considering a region of area *A*. If, at time *t*, this region contains *N*_*B*_ blood vessels and *N*_*T*_ tumour cells, then the cross-PCF, *g*_*BT*_ (*r*), is given by:

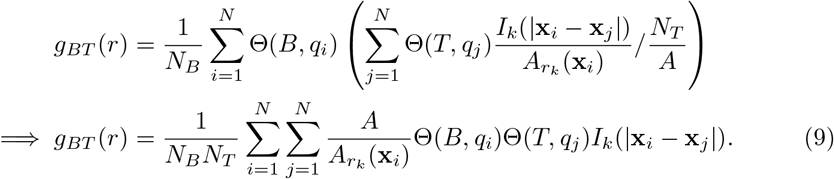

where *r* ∈ [*r*_*k*_, *r*_*k*_ + *dr*) and *N* is the total number of objects in the simulation. For each blood vessel, the cross-PCF compares the density of tumour cells in the annulus that surrounds it to *N*_*T*_ /*A*, the expected density in the annulus under complete spatial randomness (CSR). Thus, *g*_*BT*_ (*r*) > 1 indicates clustering of tumour cells at distance *r* from blood vessels and *g*_*BT*_ (*r*) < 1 indicates anti-correlation, or exclusion, of tumour cells at distance *r* from blood vessels. Cross-PCFs for other pairs of categorical variables are defined similarly. We note that the cross-PCF is not necessarily symmetric (i.e., *g*_*BT*_ ≠ *g*_*T B*_ since, for any pair of points, the annulus surrounding one point may intersect with the domain boundary while the annulus surrounding the other may not).

### Weighted pair correlation function (wPCF)

We calculate the wPCF by replacing Θ(*Q, q*_*i*_) in Equation (9) with a weighting function, 0 ≤ *w*_*p*_(*P, p*_*i*_) ≤ 1, which describes how *p*_*i*_ differs from a target phenotype, *P*. Multiple functional forms could be used for the weighting function. The relationship between the wPCF and cross-PCF is explored in more detail in S3 Appendix: Derivation of PCFs and Cross-PCFs from wPCF, and in S4 Appendix: Comparison of different weighting functions we show how the choice of weighting function affects the wPCF. For simplicity, throughout this paper we use a triangular weighting function of the form:

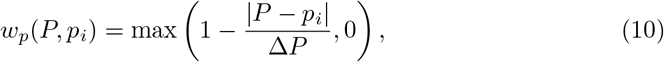

and fix Δ*P* = 0.2. Then, *w*_*p*_ ≈ 1 for cells whose phenotype *p*_*i*_ is close to the target *P* and *w*_*p*_ = 0 for those with |*P* − *p*_*i*_| > Δ*P*. We note further that *w*_*p*_(*P, p*_*i*_) → Θ(*P, p*_*i*_) as Δ*P* → 0.

Replacing Θ(*T, q*_*i*_) with *w*_*p*_(*P, p*_*i*_) in Eq (9), we define the wPCF for macrophages of target phenotype *P* and blood vessels *B*, at lengthscale *r*, as follows:

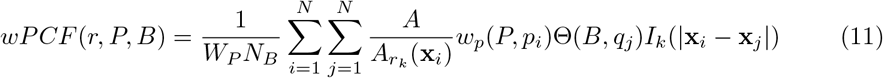

where 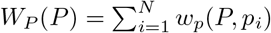 is the total ‘weight’ associated with the target label *P* across all macrophages (*W*_*P*_(*P*) replaces *N*_*T*_ in Eq (9); non-macrophages have weight *w*_*p*_ = 0). Intuitively, the wPCF extends the cross-PCF by weighting the contribution of each macrophage based on how closely its phenotype matches the target phenotype.

In Fig 3 we present two examples showing how the wPCF characterises spatial correlations between objects with a continuous label *p* (coloured circles, analogous to macrophages with phenotype *p*) and objects with a categorical label (magenta crosses, analogous to blood vessels). In both examples, 200 crosses are uniformly distributed along the line *y* = 1, and 1000 circles are randomly placed throughout a square domain of edge length 2. In Fig 3A, the label *p*_*i*_ of a circle at (*x*_*i*_, *y*_*i*_) increases linearly with distance from the line *y* = 1 − (*p*_*i*_ = |1 − *y*_*i*_|); in Fig 3B, the label increases with the square of this distance (*p*_*i*_ = |1 − *p*_*i*_|^2^). The corresponding wPCFs are shown in the middle panels of Fig 3, for a range of target labels *P* and distances *r*. For simplicity, when a wPCF is calculated over multiple values of the label *p*, we denote the resulting metric as *wPCF* (*r, p, B*). By construction, a circle at distance *r* from the nearest cross has label *P* ≈ *r* for (A) (and *P* ≈ *r*^2^ for (B)). The two wPCFs show strong clustering along these lines and exclusion at shorter distances for points with larger labels (above the dashed lines). The weaker clustering observed below the lines is explained as follows. Consider a cross at position (*x*_*j*_, 1). In (A), the largest label associated with a circle at distance *r* from this cross is *p* = *r* (if the circle is directly above the cross). Smaller labels can also be recorded, for circles at distance *r* which are offset from the cross in the y-direction. In the bottom panels of Fig 3, we plot *wPCF* (*r, P, B*) for fixed values of the target label *P*. These curves can be interpreted as cross-PCFs for points whose labels *p*_*i*_ are “close” to *P*, and show the strongest clustering at the expected values.

**Fig 3.**
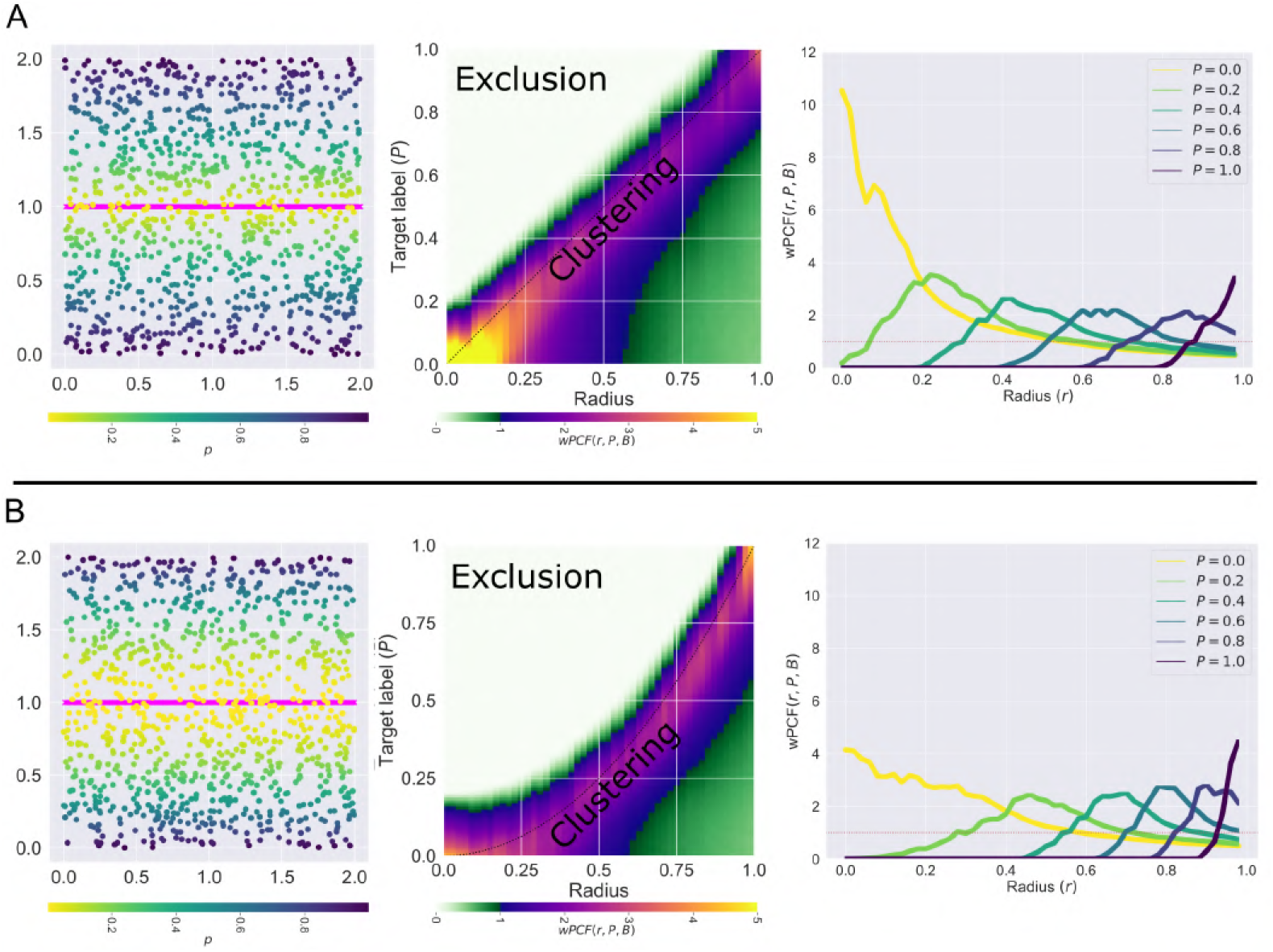
Examples for interpreting the wPCF. Two examples showing how the wPCF can identify spatial correlations between categorical objects (200 pink crosses equally spaced on the line *y* = 1) and objects with real values (1000 randomly placed circles with labels *p* ∈ [0, 1]). A: Points are labelled according to the formula *p*_*i*_ = |1 − *y*_*i*_|. B: Points are labelled according to the formula *p*_*i*_ = |1 − *y*_*i*_|^2^. Left: Point patterns consisting of equally spaced pink crosses and randomly placed circles with non-random labelling. Middle: wPCFs corresponding to the above point patterns. Dashed black lines show the lines *P* = *r* and *P* = *r*^2^, which by construction should show the strongest correlation. Right: Horizontal slices through the wPCF at fixed values of *P*. Such slices can be interpreted as a cross-PCF showing colocalisation between the pink crosses and circles with labels close to *P*.

A natural extension to the wPCF considers objects with two continuous labels, *P*_1_ and *P*_2_, say, in order to identify spatial correlations between objects with labels close to *P*_1_ and objects with labels close to *P*_2_ (e.g., colocalisation of macrophages with *p* ≈ 0 with those with *p* ≈ 1, or colocalisation of a particular macrophage phenotype with a particular concentration of CSF-1). We discuss such extensions in S5 Appendix: wPCF for comparing two continuous labels.

## Results

### Agent-based modelling generates patterns that resemble the 3 Es of immunoediting

For a given set of parameter values, we run multiple realisations of the ABM and record simulation outputs at *t* = 500 (see S2 Appendix: Tumour progression in the presence and absence of macrophages for details on simulation progression). This process generates synthetic images that resemble multiplex data, in which five categories of cells are distinguished (tumour, stroma, necrotic, vessel, macrophage) and macrophage phenotype is described using the continuous label *p*. We perform a parameter sweep of the ABM, in which we vary 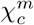, the strength of macrophage chemotaxis towards CSF-1, and *c*_1/2_, the concentration of CSF-1 at which macrophage extravasation is half-maximal. We consider 9 values of each parameter, evenly spaced for 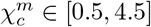 and *c*_1/2_ ∈ [0.1, 0.9]. All other parameters are held fixed at their default values (see S1 Appendix: Model Description). In Fig 4 we present typical simulation outputs at *t* = 500 for different parameter combinations (some parameter combinations are omitted to facilitate visualisation). These results show that varying 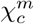 and *c*_1/2_ can generate a range of qualitative behaviours that mirror the three stages of cancer immunoediting [49]. We summarise these behaviours as follows:

- **Equilibrium**: a compact tumour mass, with macrophages confined to the surrounding stroma. The dominant macrophage phenotype is M_1_. Tumour growth is constrained, with tumour cells restricted to the mass and prevented from migrating to the vasculature by macrophage surveillance (blue box in Fig 4).
- **Escape**: the tumour has a diffuse, fragmented structure. Perivascular niches containing M_2_ macrophages and tumour cells surround blood vessels. The bulk of the tumour is infiltrated with M_1_ and transitioning macrophages, with central tumour necrosis caused by macrophages killing tumour cells (orange box in Fig 4).
- **Elimination**: total, or near-total, tumour cell elimination. Some macrophages cluster around any surviving tumour cells and the dominant macrophage phenotype is M_1_ (green box in Fig 4).

**Fig 4.**
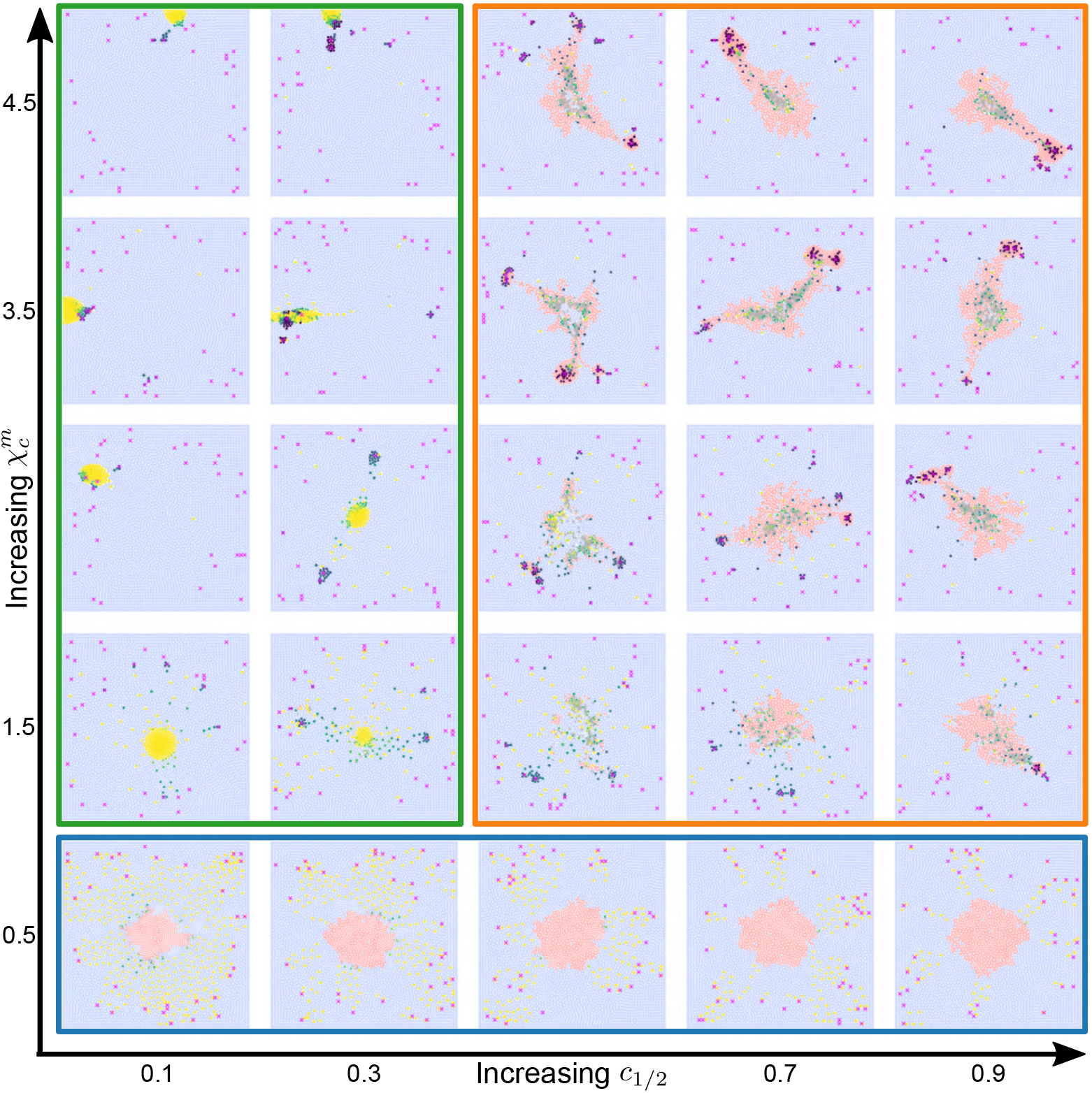
Varying macrophage sensitivity to environmental cues generates diverse patterns of tumour growth. Representative simulation endpoints for combinations of 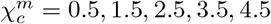 and *c*_1/2_ = 0.1, 0.3, 0.5, 0.7, 0.9. We group these into three qualitatively similar behaviours: Equilibrium - blue box: compact tumour mass, with predominantly M_1_ macrophages confined to the stroma. Escape - orange box: establishment of perivascular niches containing M_2_ macrophages, tumour cells and blood vessels. Tumour masses are asymmetrical. Elimination - green box: total or near total tumour elimination.

Equilibrium (blue box) arises for low values of 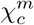 (e.g, 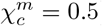). Large numbers of macrophages extravasate in response to CSF-1 but, since they are not strongly attracted to the tumour mass, they remain in the stroma. As a result, tumour growth is constrained by the macrophages, which are predominantly M_1_. When *c*_1/2_ is also small (e.g., *c*_1/2_ ⪅ 0.3), the rate of macrophage extravasation is high, and some macrophages reach the tumour boundary through random exploration of the tissue. These macrophages kill tumour cells on contact, causing the tumour mass to decrease in size.

Escape (orange box) occurs when macrophages migrate to the tumour so slowly that they do not overwhelm it. Further, exposure to TGF-*β* causes the macrophages to transition to an M_2_ phenotype. The M_2_ macrophages migrate towards nearby blood vessels, up spatial gradients in CXCL12, and the CSF-1/EGF paracrine loop enables tumour cells to trail behind them. If these tumour cells reach the vasculature, we assume that tumour cells enter the vasculature and metastasise to other parts of the body. Therefore, we denote such simulations as tumour escape.

Elimination (green box) occurs when the rate of macrophage extravasation is very high (low values of *c*_1/2_). Tumour elimination occurs because the M_1_ macrophages are strongly attracted to the tumour mass and kill tumour cells before they are ‘reprogrammed’ to an M_2_ phenotype. Strong chemotactic sensitivity to CSF-1 (large values of 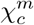) can cause the macrophages to cluster around the last tumour cells to be eliminated.

### The wPCF clarifies the relationship between macrophage phenotype and spatial distribution

Fig 5 illustrates how resolving macrophage phenotype as a continuous variable enhances interpretation of the spatial patterns that macrophages adopt in solid tumours.

**Fig 5.**
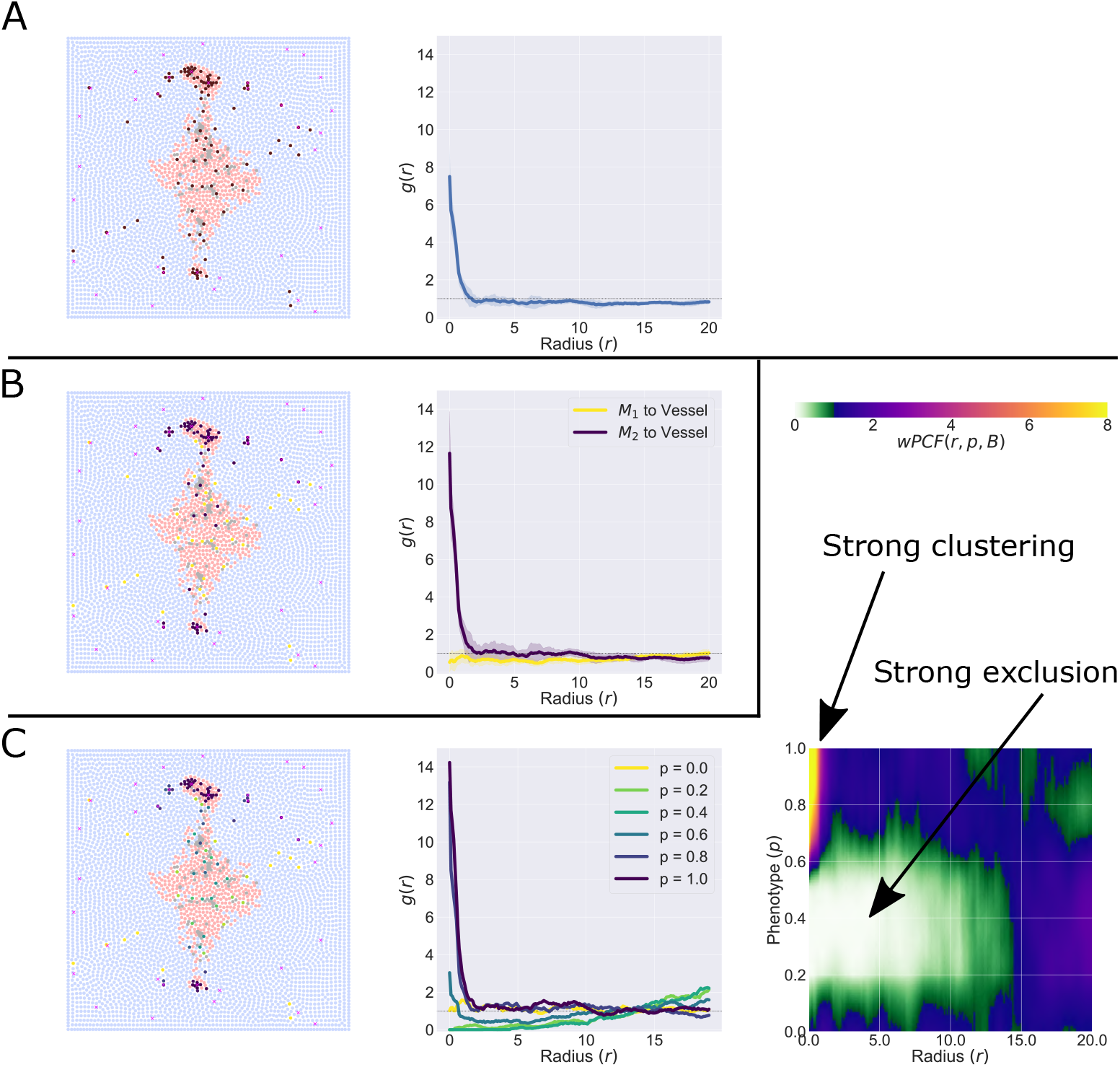
wPCF shows how macrophage phenotype and spatial distribution are related. A: treating macrophages as a single population shows clustering between macrophages and tumour cells in the cross-PCF. B: Defining two populations of macrophages (M_1_, with *p* ≤ 0.5, or M_2_ with *p* > 0.5) shows differences in spatial localisation: M_1_ macrophages are randomly spread through the domain, while M_2_ macrophages are clustered around blood vessels. C: Using the full phenotype spectrum reveals three macrophage subpopulations, which can be distinctly seen in the wPCF. Macrophages with *p* ≈ 0 have no significant spatial relationship with blood vessels, macrophages with 0.6 ⪅ *p* have strong short range colocalisation with blood vessels, and macrophages with 0.1 ⪅ *p* ⪅ 0.6 are strongly excluded from blood vessels at distances up to 15 cell diameters. As well as the full wPCF (right), we show cross-sections of the wPCF which illustrate that the wPCF has the same interpretation as the cross-PCFs in A and B, but with more resolution in macrophage phenotype.

In Fig 5A, macrophage phenotype is not resolved. The cross-PCF between macrophages and blood vessels reveals strong short-range clustering. In Fig 5B the macrophages are partitioned into two subpopulations: without loss of generality, M_1_ macrophages have *p* ≤ 0.5 while M_2_ macrophages have *p* > 0.5. We calculate the cross-PCF between each macrophage subpopulation and blood vessels. The resulting cross-PCFs show that M_2_ macrophages are strongly clustered around blood vessels, while the M_1_ macrophages are not significantly associated with the blood vessels at any length scale.

In Fig 5C macrophage phenotype is viewed as a continuous variable and we compute the wPCF between the macrophages and the blood vessels (*wPCF* (*r, p, B*)). The wPCF identifies three distinct macrophage populations, rather than the two populations used in Fig 5B. The spatial positions of macrophages with *p* ≈ 0 and blood vessels are not strongly correlated, as for the M_1_ population in Fig 5A. Macrophages with 0.6 ⪅ *p* exhibit strong short range colocalisation with blood vessels, as for the M_2_ population in Fig 5B. The wPCF identifies a third population of macrophages with 0.1 ⪅ *p* ⪅ 0.6 which is strongly excluded from blood vessels at distances up to approximately 15 cell diameters. This distance corresponds to the approximate distance from blood vessels to the tumour core, suggesting that these macrophages are localised inside the tumour mass.

### wPCFs produce signatures that distinguish the 3 Es of immunoediting

In Fig 6 we analyse the spatial patterns generated by the ABM for different values of 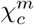 and *c*_1/2_. These patterns resemble Equilibrium (A, 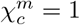, *c*_1/2_ = 0.8), Escape (B, 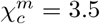, *c*_1/2_ = 0.7) and Elimination (C, 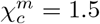, *c*_1/2_ = 0.3). For each simulation, we compute *wPCF* (*r, p, B*), *wPCF* (*r, p, T*) and *g*_*BT*_ (*r*), to characterise the pairwise spatial relationships between macrophages of different phenotypes, blood vessels, and tumour cells. We define the combination of these three statistics as a ‘PCF signature’ for our simulations.

**Fig 6.**
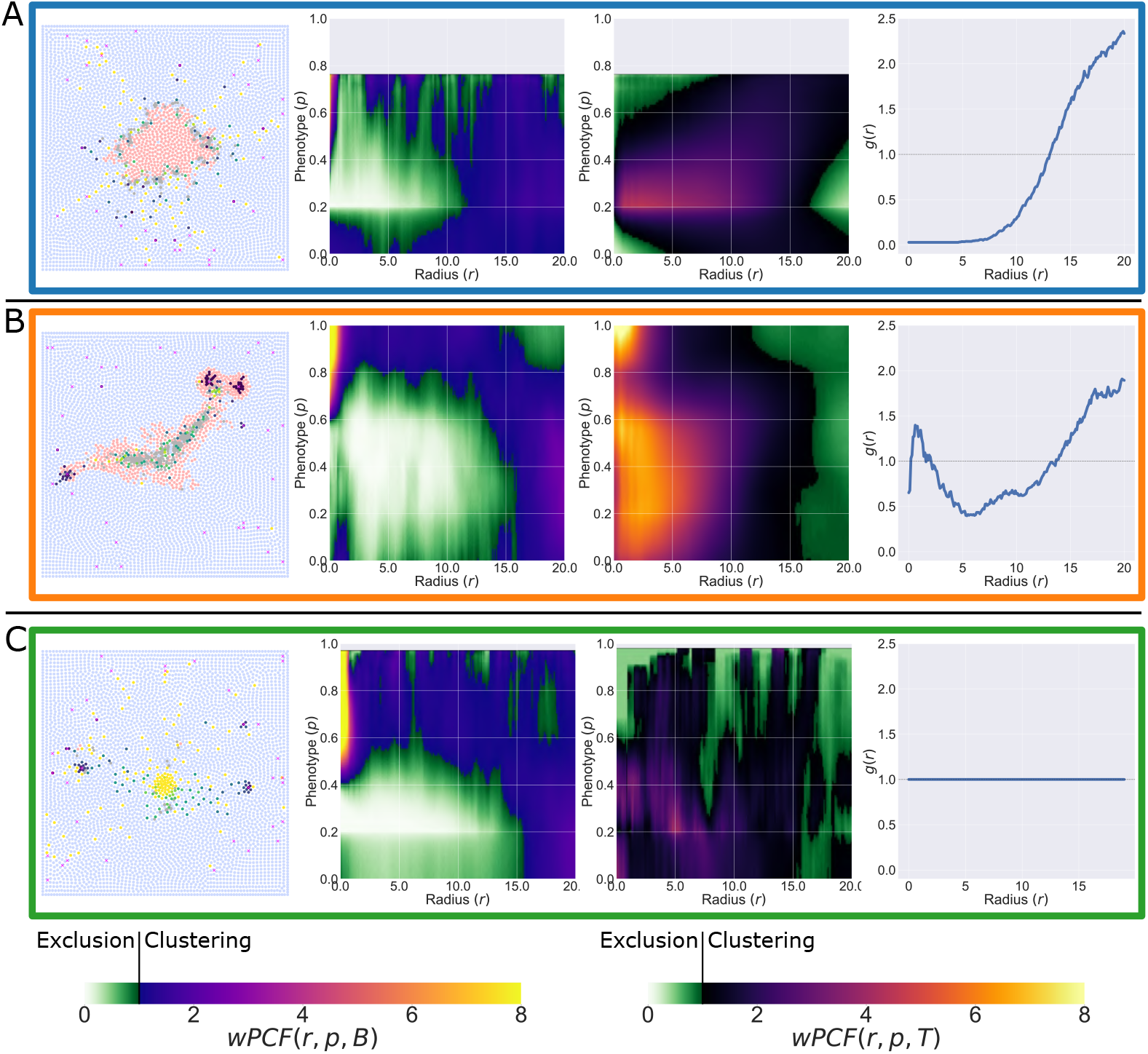
‘PCF signatures’ for Equilibrium, Escape and Elimination. We consider parameter combinations representing Equilibrium (A, 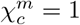, *c*_1/2_ = 0.8), Escape (B, 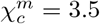, *c*_1/2_ = 0.7) and Elimination (C, 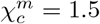, *c*_1/2_ = 0.3). For each, we show a representative simulation at *t* = 500. The wPCFs describing macrophage relationships with blood vessels and macrophage relationships with tumour cells are shown, alongside the cross-PCF describing blood vessel to tumour cell relationships (each averaged over 10 simulation repetitions).

Equilibrium simulations (Fig 6A) may not contain all macrophage phenotypes. Therefore, their wPCFs may be undefined for some values of *p* (in this case, for 0.78 ⪅ *p*.) There is a marked difference in the spatial localisation of macrophages with *p* ≈ 0 and those with *p* > 0.2, with mid-phenotype macrophages exhibiting short-range exclusion from blood vessels but not from tumour cells. In this case, macrophages which have not been exposed to TGF-*β* (*p* = 0) are restricted to the stroma while those with larger phenotype values cluster around the tumour mass, at distance from the blood vessels. The cross-PCF between blood vessels and tumour cells, *g*_*BT*_ (*r*), indicates strong exclusion between blood vessels and tumour cells at distances of up to approximately 12.5 cell diameters, which is comparable to the exclusion distance between macrophages of intermediate phenotype and blood vessels.

For the Escape simulations (Fig 6B), macrophages with *p* = 1 cluster (*r* ≈ 0) with blood vessels and tumour cells. With the peak in *g*_*BT*_ (*r*) near *r* = 0, this indicates the formation of perivascular niches containing tumour cells, blood vessels and M_2_ macrophages, as reported by Arwert *et al*. [33]. Macrophages with an intermediate phenotype are strongly excluded from blood vessels at radii up to 15 cell diameters and strongly associated with tumour cells at radii up to approximately 10 cell diameters. Taken together with the exclusion of tumour cells from blood vessels indicated by *g*_*BT*_ (*r*) for 2.5 ⪅ *r* ⪅ 12.5, this is characteristic of a central tumour mass populated with transitioning macrophages and the formation of perivascular niches.

Finally, for Elimination simulations (Fig 6C), there are no strong correlations between tumour cells and macrophages and *wPCF* (*r, p, T*) is extremely noisy (because there are very few tumour cells). Similarly, *g*_*BT*_ (*r*) ≈ 1, since most simulations with this parameter set have no tumour cells. *wPCF* (*r, p, B*) is similar to that shown in Fig 6B, indicating that the macrophage distribution for Elimination is similar to that for Escape (M_2_ macrophages cluster around blood vessels, and transitioning macrophages localise in the domain centre, at distance from the vasculature). This further suggests that the time courses for Elimination and Escape simulations may be similar at early times.

### Dimension reduction via PCA permits quantitative comparison of PCF signatures

The parameter sweep described in Fig 4 contained 737 individual simulations, with 7-10 stochastic realisations conducted for each parameter combination (limited by availability of HPC resources). Each image was manually classified as Equilibrium, Escape or Elimination according to the most prominent behaviour displayed (see inset of Fig 7 for parameter combinations and labels). We allocated 371 of these images into a training dataset (simulations with ‘iteration number’ ∈ [0, 4]) and the remaining 366 into a testing dataset (simulations with ‘iteration number’ ∈ [5, 9]). For each image, we computed *wPCF* (*r, p, B*), *wPCF* (*r, p, T*) and *g*_*BT*_ (*r*) to form the PCF signature described in the previous section. We then vectorised and concatenated the three spatial statistics, to form a high-dimensional vector (38,773 entries).

**Fig 7.**
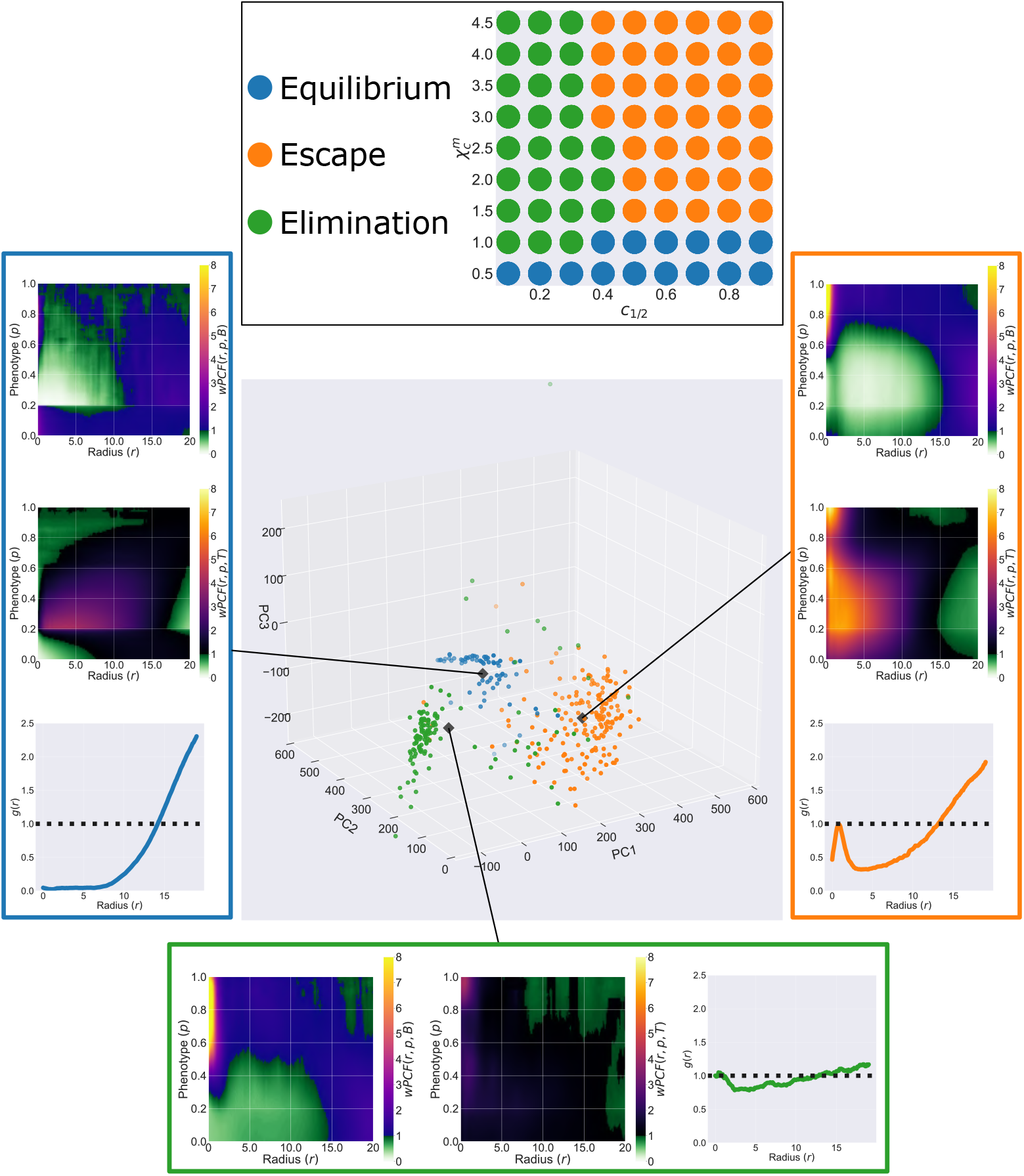
Projecting PCF signatures onto their first three principle components resolves the 3 Es of immunoediting. Top: labels assigned to each parameter combination at *t* = 500. Labels are manually assigned based on the predominant behaviour observed across all realisations of that parameter set. Main: Projection of the vectorised PCF signatures from outputs at *t* = 500 of 371 ABM simulations onto their first three principal components. Simulations cluster according to their label (manually defined as Escape, Elimination or Equilibrium as per the inset above). The top 100 principal components for the centroid of each cluster have been converted back into wPCF and cross-PCF signatures, and the corresponding *wPCF* (*r, p, B*), *wPCF* (*r, p, T*) and *g*_*BT*_ (*r*) to each are shown. Each inset has the same interpretation as the PCF signatures in Fig 6, showing that conversion between PCF signatures and PCA-space is straightforward.

We then applied dimensionality reduction to the training dataset. In Fig 7, we use principal component analysis (PCA) to project these high-dimensional vectors onto their first three principal components (the first principal component lies in the direction that maximises the variance in the data; each successive principal component maximizes the remaining variance in the data and is orthogonal to all previous principal components). Projecting the data onto the first 3 principal components shows that the synthetic images cluster according to their labels, suggesting that *wPCF* (*r, p, B*), *wPCF* (*r, p, T*) and *g*_*BT*_ (*r*) capture sufficient information to distinguish between the three qualitative behaviours that the ABM exhibits (i.e., the three Es of immunoediting).

The wPCFs and PCFs associated with the centroids of each cluster are presented in the inset in Fig 7, and are obtained by summing the first 100 principal components that define each centroid. These PCF signatures are consistent with those presented in Fig 6, and suggest that the first three principal components associated with the simulation output from the ABM at *t* = 500 could be used to classify it as Escape, Elimination or Equilibrium.

The 371 data points shown in Fig 7 were then used as training data for a basic support vector machine (SVM) with a radial basis function kernel, based on the first 100 principal components of the PCF signatures. We used the SVM to predict the labels of a testing set of the 366 simulations in the test dataset, and obtained 91.5% accuracy.

We applied the classifier to simulation outputs from a second parameter sweep, this time randomly choosing values of the same two parameters as previously (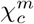 and *c*_1/2_) alongside randomly choosing four additional parameters (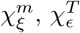, *c*_1/2_ and *g*_*crit*_) in order to generate a wider potential variety of simulation outcomes. This produced 431 additional images, which were manually labelled as Equilibrium, Escape or Elimination at *t* = 500. Table 1 shows the performance of the classifier at predicting the labels of these simulations. The classifier identifies Escape simulations with extremely high accuracy, correctly identifying 97.2% of simulations in which tumour intravasation and metastasis is present.

**Table 1.** Classification performance for 100 principal components. Classifier accuracy (number of classifications and percentage of true classifications assigned to that class) for SVMs trained on the first 100 principal components, based on 431 manually labelled simulations at *t* = 500 with parameters randomly sampled within a 6-parameter space. Bold fields show correct classifications.

|  |  | Predicted classification |  |  |
| --- | --- | --- | --- | --- |
|  |  | Equilibrium | Escape | Elimination |
| True classification | Equilibrium | **78 (65%)** | 26 (22%) | 16 (13%) |
|  | Escape | 0 (0%) | **104 (97%)** | 3 (3%) |
|  | Elimination | 18 (9%) | 14 (7%) | **172 (84%)** |

### ABM simulations may transition between the three Es of immunoediting over time

We have shown that ABM data from a single timepoint can be classified as Equilibrium, Escape or Elimination based on their PCF signatures. Since the ABM simulations are dynamic, we can use the methods used to create Fig 7 to investigate how the qualitative behaviour of an ABM simulation changes over time.

The results presented in Fig 8 derive from an ABM simulation with 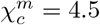 and *c*_1/2_ = 0.3, which we classify as ‘Elimination’ based on its output at *t* = 500. The time series in Fig 8 show how, as the tumour develops, the simulation transitions from ‘Equilibrium’ (compact mass, *t* = 250) through to ‘Escape’ (*t* = 350, 400) and, ultimately, to ‘Elimination’.

**Fig 8.**
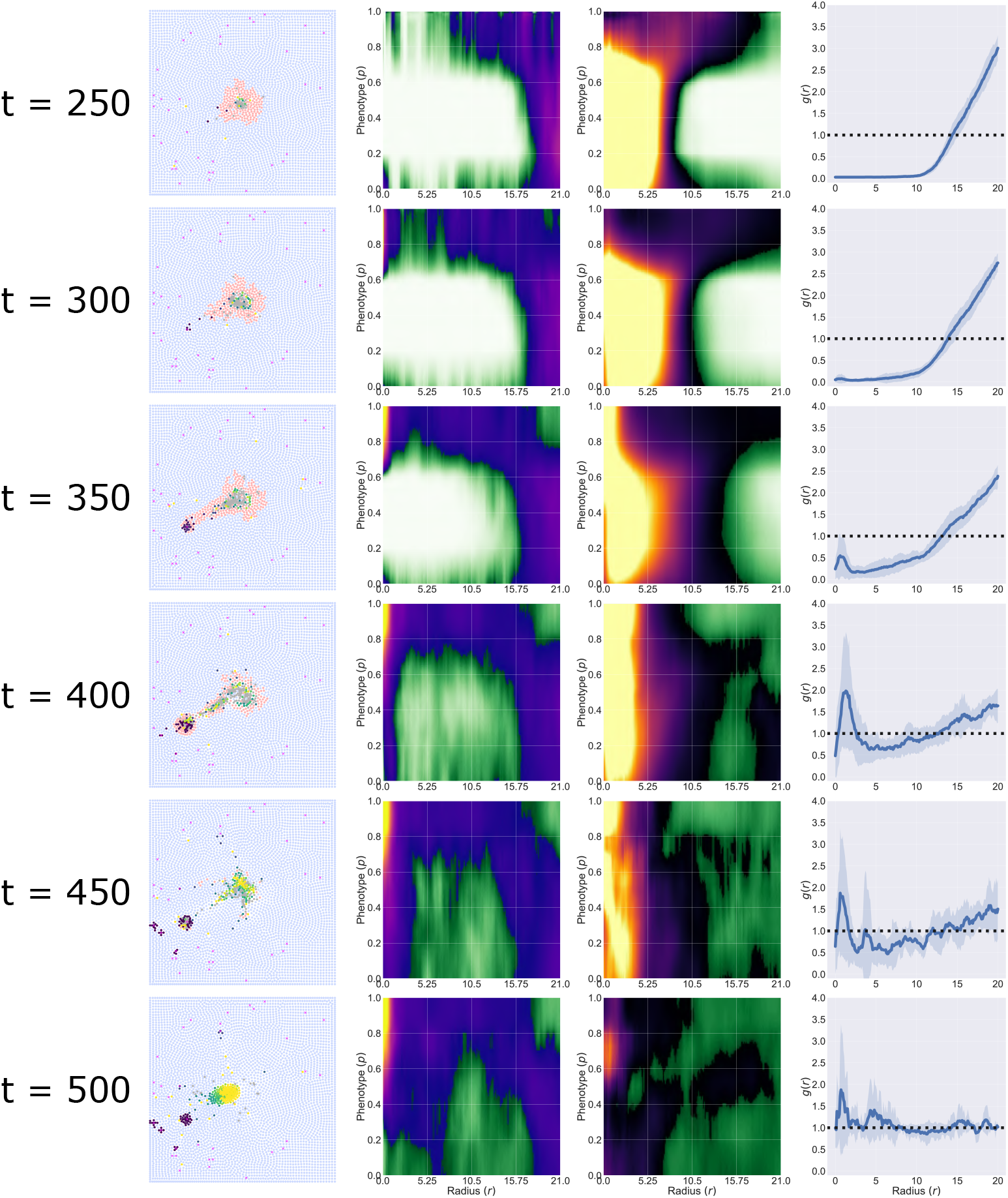
Dynamic evolution of an ABM simulation. Time series showing the evolution of an ABM simulation generated with 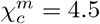 and *c*_1/2_ = 0.3, and all other parameter values fixed at the default values listed in S1 Appendix: Model Description. At different timepoints, the simulation exhibits behaviours which are consistent with Equilibrium, Escape and Elimination. The corresponding PCF signatures show how the ABM progresses through these stages.

We calculate PCF signatures for this ABM simulation every 10 hours, and project them onto the first three principal components, using the process in Fig 7. The resulting trajectory is depicted in Fig 9, with points coloured according to their time and the insets showing the corresponding synthetic images.

**Fig 9.**
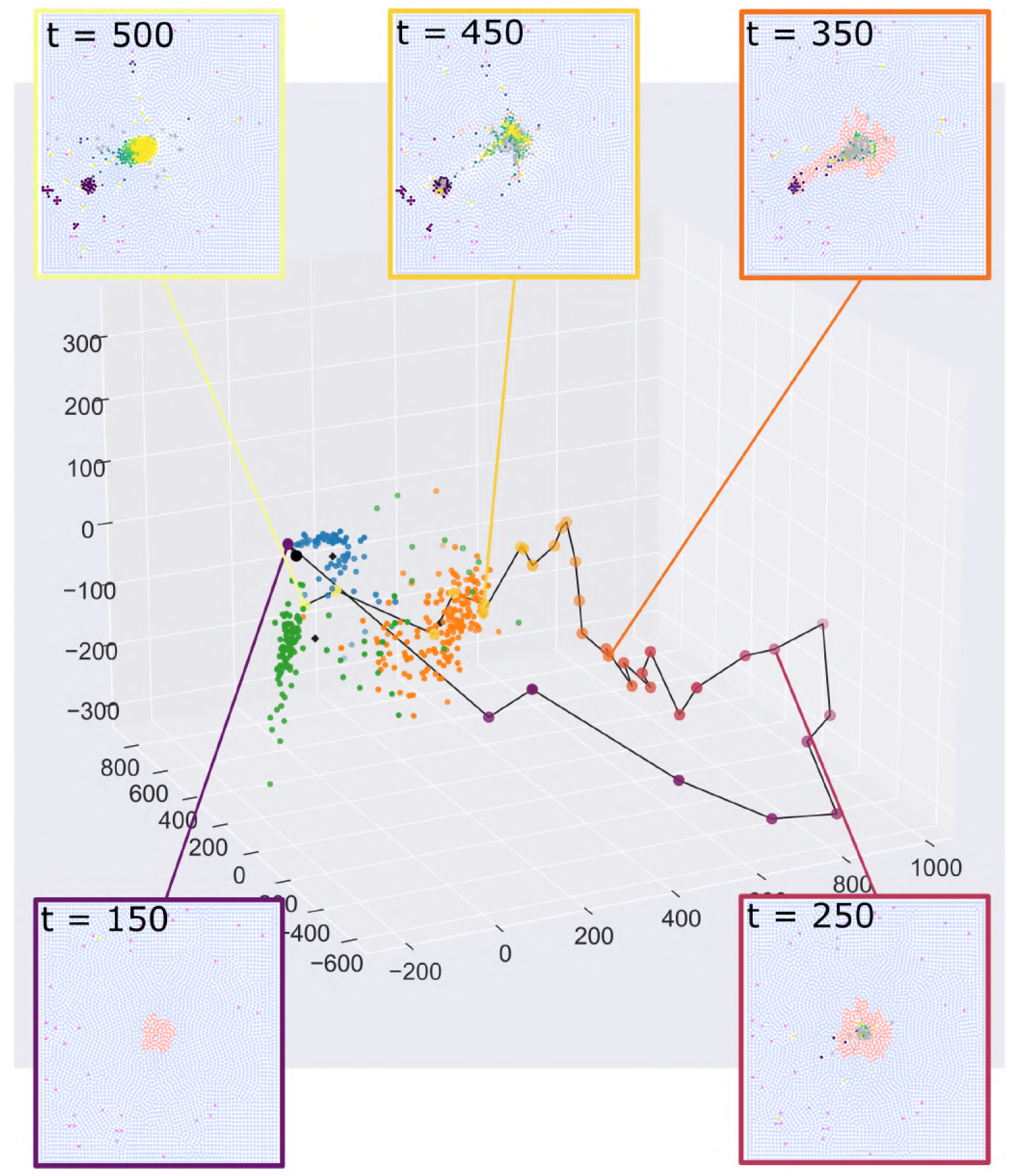
Dynamic evolution of an ABM simulation through PCA space. A single simulation can exhibit Equilibrium (*t* = 150), Escape (*t* = 350, 450) or Elimination (*t* = 500) at different times. These changes are captured by the movement of the PCF signature through PCA space.

At early times, the images localise within the Equilibrium cluster (blue cluster; *t* = 150). Once macrophages start to appear in the images, the trajectory moves away from the three clusters, due to noise in the wPCFs caused by the small number of macrophages relative to the *t* = 500 simulations which define the clusters (*t* = 250). As the number of macrophages increases, the trajectory moves closer to the Escape cluster (orange cluster; *t* = 350 and *t* = 450). Finally, as the tumour cells are killed and removed from the simulation, the trajectory moves into the Elimination cluster (green cluster; *t* = 500).

This study highlights two related challenges. First, spatial data from a single timepoint may not be predictive of past or future behaviour: an ABM may exhibit multiple behaviours at different timepoints. Second, for this simulation, some tumour cells successfully migrate to neighbouring blood vessels prior to the tumour’s elimination. In practice, these tumour cells could enter the vasculature and spread to other parts of the body. Given information about the tumour’s spatial composition at *t* = 500 only, we will classify this image as Elimination without identifying the Escape behaviour present at earlier times.

## Discussion

In this paper we have introduced a new spatial statistic, the wPCF, which extends the cross-PCF to point clouds labelled with a mixture of categorical and continuous labels. We demonstrated its utility by applying it to synthetic data generated from a new ABM that simulates macrophage interactions with a tumour growing in a 2D vascular tissue.

Blood vessels and tumour cells are categorically labelled, while macrophages have a continuous phenotypic label. The wPCF reveals spatial correlations between the different cell types.

The ABM focusses on the impact that phenotypic heterogeneity in the macrophage population has on the tumour’s patterns of growth. By varying parameters that control macrophage sensitivity to environmental cues, we used the ABM to generate a range of synthetic data that spans the three Es of immunoediting (Equilibrium, Escape or Elimination). We showed that wPCFs and cross-PCFs can be combined to produce a high dimensional ‘PCF signature’ which characterises the relative locations of macrophages, tumour cells and blood vessels. Our results suggest that, given suitable training data, the PCF signature could be used to automate the classification of images or point patterns into different clusters, by using PCA to project the PCF signature onto a lower dimensional space.

One advantage of testing the wPCF on synthetic data from an ABM is its ability to generate time-series data. Our analysis of dynamic output from an ABM simulation showed how the PCF signature may vary over time. A given simulation may transition between different states, suggesting that caution is needed when using a single time point to make predictions about a tumour’s future growth and response to treatment. This study also highlights one of the benefits of developing mathematical models of a biological system: the ABM can be used to investigate questions that would be challenging to address with existing experimental techniques. For example, it is not currently feasible to apply multiplex imaging to the same tissue at multiple timepoints.

There are many directions for extending and improving the ABM. For example, including tumour cell intravasation would prevent situations in which an Escape signature can progress to Elimination at a later timepoint, as in Fig 8. In practice, tumour cells would have entered the vasculature and may have established metastases. In future work, we will also explore ABM extensions which incorporate therapies, such as radiotherapy or immunotherapy, and investigate whether interactions between different cell types are predictive of response (see, e.g., [64–66]). We will consider whether the spatial distributions of cells at different timepoints must be accounted for when making such predictions. We could further extend the ABM to account for multiple immune cell subtypes, such as T cells, or stromal cells such as fibroblasts [67].

The robustness of the wPCF should be tested through application to other types of ABMs that simulate tumour growth and generate similar outputs. In addition to the Chaste framework used here [51–53], other candidate ABM frameworks include PhysiCell [81], HAL (Hybrid Automata Library) [82], and CompuCell3D [83, 84].

In this paper we used the wPCF and cross-PCF to describe and classify synthetic data generated by an ABM. However, the wPCF has application to a wider range of scientific fields. In particular, the wPCF can account for continuous variation in expression levels of cellular markers which characterise multiplex imaging modalities, such as multiplex immunohistochemistry or Imaging Mass Cytometry. Equally, the wPCF could be used to describe any structured cell population, for example continuous labels describing the stemness or differentiation status of cancer cells, the exhaustion level of T-cells, or the oxygen concentration experienced by cells within a tissue sample. Beyond biology, the wPCF could be used to analyse data from other applications in which PCFs have proven useful, including astronomy [19, 20] and ecology [16–18]. The wPCF can be extended to compare the spatial distributions of point patterns with multiple continuous labels. In Fig 1 expression levels of CD68, CD163, and CD204 were used to define distinct macrophage subtypes; each marker could represent a separate continuous structure label. We show how the wPCF can be used to describe the spatial distribution of points with two continuous labels in S5 Appendix: wPCF for comparing two continuous labels.

Future work will involve applying the wPCF to multiplex imaging data, in order to validate its use in biological and clinical settings. Applying such statistics to medical images would enable their high-throughput, automated quantification and comparison in a manner that goes beyond expert visual inspection and is more interpretable than AI approaches [57, 58]. We note also that while in this paper we focus on correlation functions, alternative metrics, including topological data analysis, can describe spatial features such as immune deserts that exist in noisy data [30], or changes in tumour and vascular architecture in response to radiotherapy [59]. Multiple spatial statistics can be combined to obtain more detailed descriptions of 2D data [60], or new statistics can be derived from networks of cell contact [61] or observations of immune cell locations [62, 63].

In this paper we presented a proof-of-concept SVM classifier to show how multiple statistics can be combined to classify data, with PCA acting as a dimension reduction technique which permits a classifier to be trained on high dimensional statistics without sacrificing their interpretability (as is generally required for AI approaches). While this approach works acceptably, there is considerable scope for optimisation around i) the choice of statistics, ii) the choice of classification method, and iii) the choice of dimension reduction tool. In future work, we will explore alternative choices at each of these stages in order to better optimise a pipeline for classification of point clouds based on their spatial structure. We will also use a wider range of techniques, such as topological data analysis, to further characterise the outputs from different simulations and timepoints.

This paper demonstrates an exciting proof-of-concept: statistics which describe different aspects of cell localisation can be combined to classify, describe and analyse synthetic and biological point clouds.

## Supporting information

## S1 Appendix: Model Description

Our ABM extends an existing ABM that describes infiltration of microbeads [50] and macrophages [30] into tumour spheroids growing *in vitro*. Here, we consider an *in vivo* scenario, where tumour cells are embedded within a tissue containing stromal cells and oxygen is supplied by blood vessels. We use an overlapping-spheres model, representing each cell as a point whose movement and position are determined by balancing the forces that act on it. We distinguish four types of agent: **macrophages**, **tumour cells**, **stromal cells** and **necrotic cells**. We also consider five diffusible species: **oxygen** (*ω*(**x**, *t*)), **colony stimulating factor-1** (CSF-1, *c*(**x**, *t*)), **transforming growth factor-***β* (TGF-*β*, *g*(**x**, *t*)), **epidermal growth factor** (EGF, *ϵ*(**x**, *t*)), and **C-X-C motif chemokine 12** (CXCL12, *ξ*(**x**, *t*)). Their dynamics are defined by reaction-diffusion equations (RDEs). Following [30, 50], we use the open source Chaste (Cancer, Heart and Soft Tissue Environment) modelling environment to generate simulation [51–53].

## Diffusible species

The distribution of the diffusible species is modelled using RDEs, with each equation solved numerically on a triangular finite element mesh that spans the domain. We consider a square domain Ω, with height and width equal to 50 cell diameters (approximately 1 mm in dimensional units). In this Section we describe the role of each diffusible species, the factors that govern their rates of production and depletion, and the RDEs that define their evolution. Fig 2 in the main text summarises how the diffusible species interact with tumour cells and macrophages. Since we assume oxygen and CXCL12 are produced from the (static) blood vessels, we make a quasi-steady state assumption for their distribution. By contrast, changes in the distributions of EGF, TGF-*β* and CSF-1 are assumed to happen on the slower timescale associated with tumour cell and macrophage movement. Therefore we do not make the quasi-steady assumption when calculating these distributions.

## Oxygen (*ω*)

Oxygen is supplied by blood vessels which are represented by static point sources, and is consumed by tumour cells and stromal cells as it diffuses through the domain. We assume that the oxygen concentration at each blood vessel is constant, *ω*_0_.

Since the timescale of diffusion for an oxygen molecule is faster than the timesteps used in our model to describe cell movement, we assume that the distribution of oxygen can be approximated using a steady state solution [30, 50]. For simplicity, we rescale the oxygen concentrations with a factor of *ω*_0_, so that *ω* = 1 at each blood vessel. The governing equation is then given by

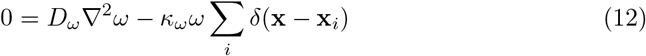

for **x** ∈ Ω, where **x**_*i*_ is the location of stromal or tumour cell *i*, the parameter *D*_*ω*_ is the assumed constant diffusion coefficient of oxygen, *κ*_*ω*_ is the oxygen consumption rate and *δ* is the delta function (*δ*(**x**) = 1 when **x** = 0, *δ*(**x**) = 0 otherwise). We impose Neumann boundary conditions 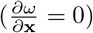 on domain boundaries, and assume that initially *ω*(**x**, *t* = 0) = 1 for all **x** that do not represent a blood vessel.

## CXCL12 (C-X-C motif chemokine 12, *ξ*)

CXCL12 is produced by perivascular cancer-associated fibroblasts (CAFs) and acts as a diffusible chemoattractant for M_2_ macrophages [33]. We suppose that CAFs localise close to blood vessels, and, hence, make the simplifying assumption that blood vessels act as constant sources of CXCL12. We therefore assume that the concentration of CXCL12, *ξ*, is maintained at a fixed value *ξ* = *ξ*_0_ at all blood vessels. We assume that CXCL12 decays naturally at a constant rate, *λ*_*ξ*_. The distribution of CXCL12 in the domain is then given by

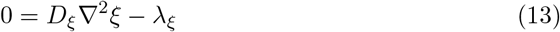

where *D*_*ξ*_is the diffusion coefficient for CXCL12. We impose Neumann boundary conditions 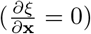 on domain boundaries, and assume that initially *ξ*(**x**, *t* = 0) = 0 for all **x** that do not represent a blood vessel.

## CSF-1 (Colony stimulating factor-1, *c*)

The macrophage chemoattractant CSF-1 is produced by tumour cells. It acts with EGF as part of a paracrine loop [68, 69] and stimulates macrophage extravasation [33]. We assume that CSF-1 is produced at a constant rate, *κ*_*c*_, by each tumour cell, and decays at a constant rate, *λ*_*c*_. The distribution of CSF-1 is therefore described by the equation

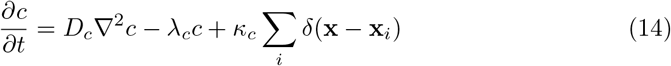

where *D*_*c*_ is the diffusion coefficient of CSF-1 and the sum runs over all tumour cells *i*. We impose Neumann boundary conditions 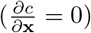 on domain boundaries, and assume that initially *c*(**x**, *t* = 0) = 0 for all **x**.

## TGF-*β* (Transforming growth factor-*β*, *g*)

TGF-*β* causes macrophages to alter their phenotype, and is produced by tumour cells. We suppose that it decays at a constant rate, *λ*_*g*_, and is produced by tumour cells at a constant rate, *κ*_*g*_. Combining these effects, we arrive at the following RDE for *g*(**x**, *t*):

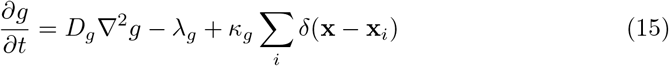

where *D*_*g*_ is the diffusion coefficient of TGF-*β* and the sum is over all tumour cells, *i*. As for CSF-1, we impose Neumann boundary conditions (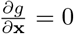 on domain boundaries), and assume that *g*(**x**, *t* = 0) = 0 ∀**x** ∈ Ω.

## EGF (Epidermal growth factor, *ϵ*)

EGF is a diffusible chemoattractant for tumour cells that is produced by tumour associated macrophages. We assume that EGF is produced by macrophages at a rate that is linearly dependent on their phenotype *p*, so that macrophages with an extreme M_1_ phenotype (*p* = 0) produce no EGF and macrophages with an extreme M_2_ phenotype (*p* = 1) produce *κ*_*ϵ*_. We assume that EGF naturally decays at a constant rate, *λ*_*ϵ*_. Combining these effects, we obtain the following RDE for EGF, *ϵ*(**x**, *t*):

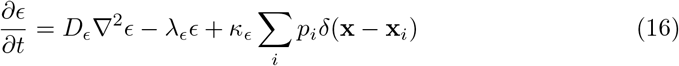

where *D*_*ϵ*_ is the constant diffusion coefficient of EGF, *κ*_*ϵ*_ is the maximum rate of production of EGF, the sum is over all macrophages *i*, and *p*_*i*_ ∈ [0, 1] is the phenotype of macrophage *i* and is defined below. As for CSF-1 and TGF-*β*, we impose Neumann boundary conditions (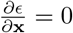 on the boundaries of the domain Ω), and assume that *ϵ*(**x**, *t* = 0) = 0 ∀**x** ∈ Ω.

## Agents

Our model distinguishes four types of agents: **macrophages**, **tumour cells**, **stromal cells** and **necrotic cells**. A fixed number of static **blood vessels** are also distributed throughout the domain. The location of each agent is represented by its cell centre.

We use Newton’s second law to determine the equations of motion for macrophages, tumour cells, stromal cells and necrotic cells. Neglecting inertial effects in the overdamped limit, the force balance for cell *i* can be written as:

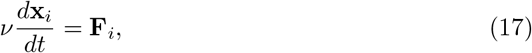

where *ν* is the drag coefficient (assuming that the drag on a cell is proportional to its velocity), and **F**_*i*_ denotes the net force acting on cell *i*. The forces that act on each cell type are indicated in Fig 2.

All cells are subject to mechanical forces due to cell-cell interactions (incorporating repulsion due to volume exclusion and attraction due to intercellular adhesion). Here we use the overlapping spheres approach [70, 71], assuming that two cells interact if the distance between their cell centres is less than a fixed distance *R*_*int*_. Specifically, for cells at locations **x**_*i*_ and **x**_*j*_, if |**x**_*i*_ − **x**_*j*_| < *R*_*int*_ then the interaction force between cells *i* and *j* is parallel to the vector (**x_*i*_** − **x_*j*_**) connecting their centres. The resting spring length between the cell centres, *s*_*i,j*_, is the sum of the equilibrium spring lengths associated with each cell (*s*_*i,j*_ = *s*_*i*_ + *s*_*j*_). For most cells *i*, the resting spring length is equal to the approximate radius of a cell (*s*_*i*_ = *R*_*Cell*_ ≡ 0.5 in non-dimensional units). For newly divided and necrotic cells, *s*_*i*_ changes over time to account for cell growth and shrinkage (see below).

The net mechanical force acting on cell *i* is the sum of all interaction forces due to cells within its interaction radius:

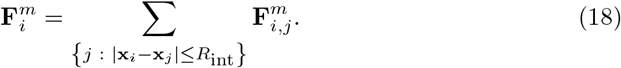

where 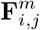 is the mechanical force between cells *i* and *j*. This force always points in the direction of the vector connecting the cell centres and has magnitude:

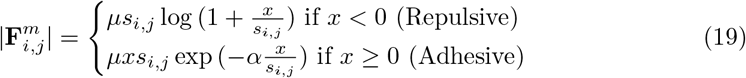

where *x* = |**x**_*i*_ − **x**_*j*_| − *s*_*i,j*_ is the overlap between cells *i* and *j*, *μ* is a parameter describing the spring stiffness and *α* determines the strength of intercellular adhesion between neighbouring cells.

We normalise lengths with the lengthscale *R*_*Cell*_, assuming that 1 cell diameter = 2*R*_*Cell*_ = 20*μ*m. Following cell division, the radius of the daughter cells is set to 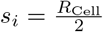 and increases linearly over one hour until *s*_*i*_ = *R*_*Cell*_. For necrotic cells, *s*_*i*_ decreases linearly to zero over 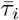 hours (see below), and then the cell is removed from the simulation. The associated spring stiffness *μ* of springs attached to a necrotic cell is reduced linearly at the same rate.

## Macrophages

## Extravasation

Macrophages extravasate from blood vessels in response to local levels of CSF-1. At each timestep, the probability that a macrophage extravasating in a one hour time period is *P*_*ex*_, where:

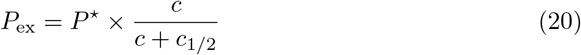

where the parameter *P*^⋆^ controls the maximum possible probability of macrophage extravasation per hour from each vessel, and *c*_1/2_ is the concentration of CSF-1 at which this probability is half-maximal.

## Phenotype

The diffusible species included in our model bind to receptors on the outer membranes of macrophages and tumour cells and regulate behaviours including chemotaxis and the production of other species. While we do not explicitly model surface receptors, the receptor CXCR4 (C-X-C motif chemokine receptor type 4) plays an important role in this system: when exposed to TGF-*β*, macrophages increase expression of CXCR4 and become sensitive to gradients of CXCL12 [33]. We model this by associating each macrophage with a phenotype, *p* ∈ [0, 1], which determines the extent to which it exhibits M_1_- or M_2_ behaviour. When a macrophage first enters the spatial domain, it has phenotype *p* = 0. Exposure to TGF-*β* causes its phenotype to increase irreversibly. The rate of change of the phenotype of macrophage *i* at location **x**_*i*_ is:

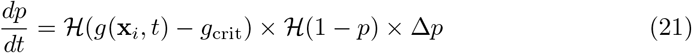

where *g*(**x**_*i*_, *t*) is the concentration of TGF-*β* at **x**_*i*_ at time *t*, 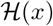 is the Heaviside function (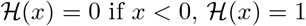 otherwise), the non-negative parameter Δ*p* determines the rate at which phenotype changes, and *g*_*crit*_ is a critical TGF-*β* threshold above which macrophage phenotype increases. When *p* = 1 the macrophage has a fully M_2_ phenotype, and *p* no longer changes.

Phenotype affects the following macrophage behaviours:

- the rate at which they produce EGF;
- the rate at which they kill tumour cells;
- their chemotactic sensitivity to spatial gradients of CSF-1 and CXCL12.

Equation (16) describes how the rate of EGF production depends on *p*; we now describe how cell killing and phenotype-dependent chemotaxis are implemented.

## Cell killing

Macrophage phenotype determines the probability that a macrophage kills a tumour cell on a given timestep. The probability that macrophage *i* of phenotype *p*_*i*_ kills a tumour cell in one hour is

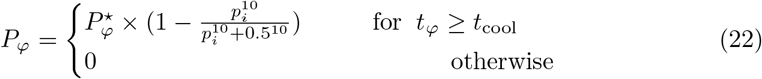

where 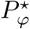 is the maximum probability that a macrophage kills a tumour cell in one hour. *t*_*φ*_ is the time since the macrophage last killed a tumour cell, and the parameter *t*_*cool*_ defines a ‘cooldown’ period during which the macrophage cannot kill another tumour cell. On each timestep the probability of each macrophage attempting cell killing is evaluated. If a macrophage attempts killing, any cells within distance 1 of the macrophage are identified and, if there is at least one tumour cell, one of the tumour cells is selected at random to be killed. The tumour cell is labelled as necrotic, and *t*_*φ*_ is reset to 0 for that macrophage.

## Force laws for macrophages

In addition to mechanical forces, macrophages are subject to forces which account for random movement and chemotactic forces due to spatial gradients of CXCL12 and CSF-1. We account for random movement via the force 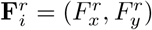 where

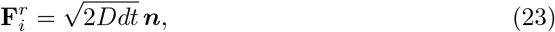

where the coefficient *D* describes the strength of the random force and ***n*** = (*n*_*x*_, *n*_*y*_), where *n*_*x*_ and *n*_*y*_ are random variables drawn from a standard normal distribution.

The net force due to chemotaxis acting on macrophage *i* depends on its phenotype, *p*_*i*_, as follows:

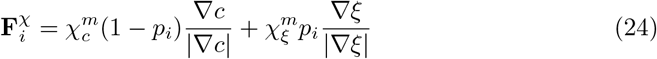

where the parameters 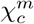 and 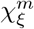 control macrophage sensitivity to spatial gradients of c ξ

CSF-1 and CXCL12 respectively. We suppose that macrophage sensitivity to CSF-1 and CXCL12 gradients scales linearly with phenotype.

The net force acting on macrophage *i* in Equation (17) is:

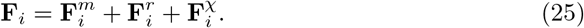

## Tumour cells

## Cell cycle and proliferation

Each tumour cell has an internal cell cycle which advances at a rate that depends on the local oxygen concentration and two oxygen thresholds, 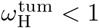 and 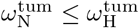. When 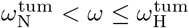, the cell becomes hypoxic and pauses its cell cycle until *ω* increases above this threshold. If 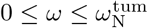, then the tumour cell dies and becomes a necrotic cell. We also account for contact inhibition in our model. If a cell’s area falls below a proportion 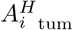 of its target area, then its cell cycle pauses until space is available for proliferation. Each tumour cell *i* has a subcellular variable denoted *T*_*i*_ which tracks its progress through the cell cycle cell as follows:

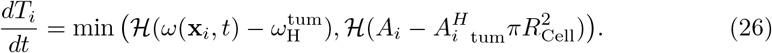

where *A*_*i*_ is the area of the cell calculated as 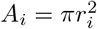 and *r*_*i*_ is the estimated cell radius calculated via the average separation between cells within the interaction radius of *i*.

Each cell has a target cell cycle duration *τ*_*i*_, drawn at random from a uniform distribution of *U* (0.75*τ*_*tum*_, 1.25*τ*_*tum*_) with average cell cycle duration *τ*_*tum*_. When *T*_*i*_ = *τ*_*i*_, cell division occurs and a new cell is placed at a distance half a cell diameter away, in a randomly chosen direction. Both cells are assigned new cell cycle durations. *T*_*i*_ is set to 0 for each cell, and then evolves according to Eq (26). The equilibrium spring length associated with the new cells is set to *s*_*i*_ = 0.5*R*_*Cell*_ to account for their reduced size, and increases linearly over the course of 1 hour until it reaches *s*_*i*_ = *R*_*Cell*_.

## Force laws for tumour cells

Tumour cell movement is governed by the force balance described in Equation (17). The net force incorporates intercellular interactions as described above, and a chemotactic force due to spatial gradients in EGF. The chemotactic force, denoted 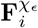, has the form

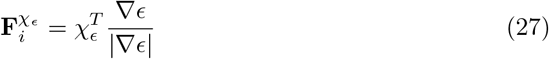

where the parameter 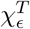 determines tumour cell sensitivity to EGF gradients. The net force used in Equation (17) is therefore

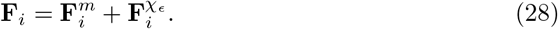

## Stromal cells

## Cell cycle and proliferation

The same cell cycle model is used for stromal and tumour cells, but it is parameterised differently to account for the increased ability of tumour cells to survive in adverse conditions such as lower oxygen environments or increased mechanical pressure from neighbouring cells. Like tumour cells, stromal cells possess a subcellular age variable *T*_*i*_ which evolves according to the equation

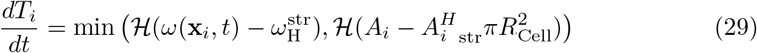

where the parameter 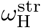 determines the oxygen threshold below which stromal cells become hypoxic and the parameter 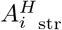 defines the proportion of a stromal cell’s target area below which the cell cycle stops.

As with tumour cells, we define a second oxygen threshold 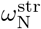, below which stromal cells become necrotic. We suppose that 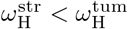 and 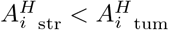 to account for the ability of tumour cells to proliferate in more adverse environments than stromal cells.

## Force laws

Stromal cells are subject to the same intercellular forces as macrophages and tumour cells, and defined by Equations (18)-(19). The force balance for stromal cells used in Equation (17) is

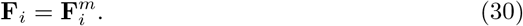

## Necrotic cells

When a stromal or tumour cell is marked for cell death, as a result of oxygen starvation or being killed by a macrophage, it irreversibly becomes necrotic. A necrotic cell *i* occupies space for 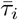 hours, where 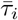 is drawn from a uniform distribution 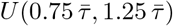 and 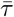 is the average duration of necrosis. Over this time period, the necrotic cell shrinks in size, by reducing its equilibrium spring constant, *s*_*i*_, at a constant rate until *s*_*i*_ = 0. The cell is then removed from the simulation. While *s*_*i*_ is being reduced, the spring constant *μ* associated with the cell is also reduced to 0 at a constant rate, to account for weakening of the intercellular forces between degrading necrotic cells and other cells.

## Initial and boundary conditions for cells

RDEs describing the diffusible species are solved numerically on a regular triangular mesh with edge length 1 cell diameter. We initialise the simulation by selecting lattice sites to act as point vessels, which do not occupy space or interact directly with cells in our model. All lattice sites more than *R*_*B*_ cell diameters from the centre of the domain are possible locations for blood vessels, and *N*_*B*_ of these sites are chosen, at random, to be blood vessels. This ensures that the centre of the domain is at least *R*_*B*_ cell diameters from the nearest blood vessel and, hence, that there is sufficient space to observe macrophage movement between blood vessels and the tumour, while also ensuring that cells near the domain boundaries are well-oxygenated.

The domain is initialised with stromal cells filling the domain in rows that are 0.75 cell diameters apart, with alternating rows offset by 0.375 cell diameters (i.e., forming a hexagonal lattice). We place four tumour cells in a cluster at the centre of the domain approximately 0.5 cell diameters apart. Stromal and tumour cells are assigned cell cycle durations *τ*_*i*_ from the relevant distributions, and cell cycle progression times *T*_*i*_ selected at random from a uniform distribution *U* (0, *τ*_*i*_). All cells are constrained to remain within the domain by imposing reflective boundary conditions.

## Schematic

The schematic in Fig 10 shows the order in which the above processes occur. After initialisation, on each timestep the concentrations of diffusible species are updated and macrophage extravasation occurs. Individual cell cycles, target spring lengths, phenotype changes, and proliferation are then updated. Finally, cells are moved to their new locations.

**Fig 10.**
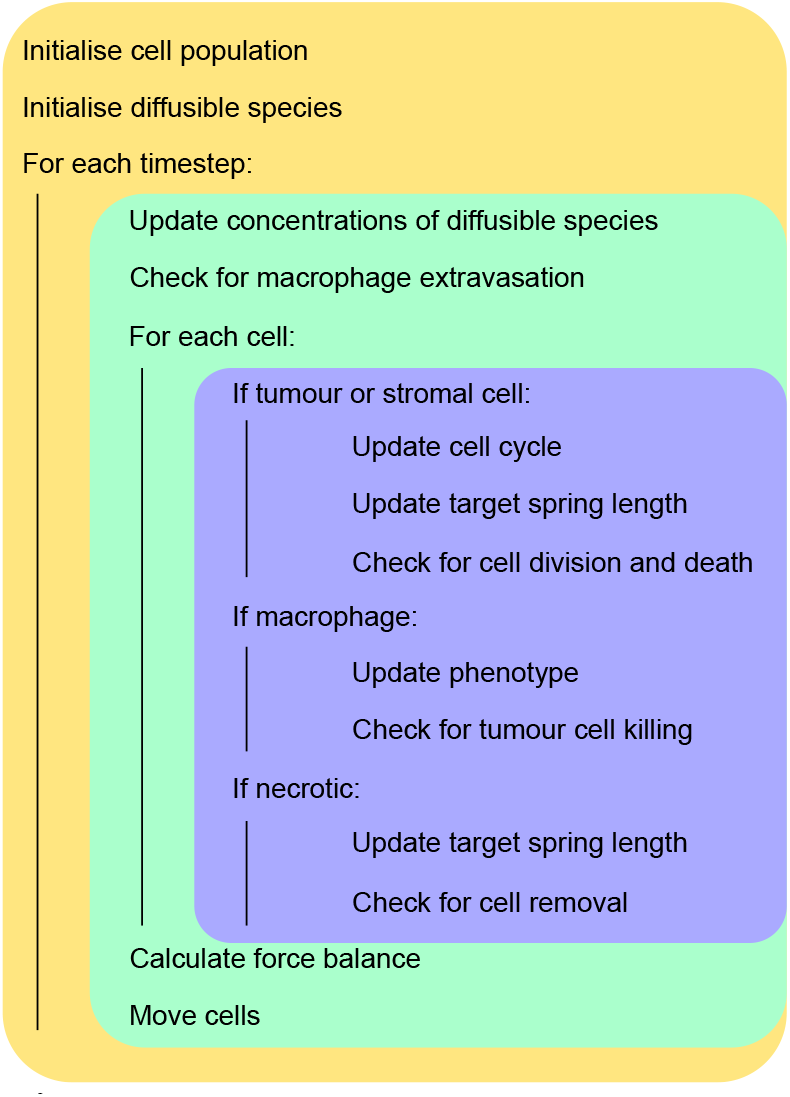
Model overview. Schematic showing an overview of the order in which the model is initialised, updated at each timestep, and updated at a cell level.

## Table of ABM parameters

In Table 2 we list the model parameters, their default dimensional and dimensionless values or ranges, and supporting references where these are available. Values of some parameters, indicated with *, have been estimated based on model behaviour.

**Table 2.**
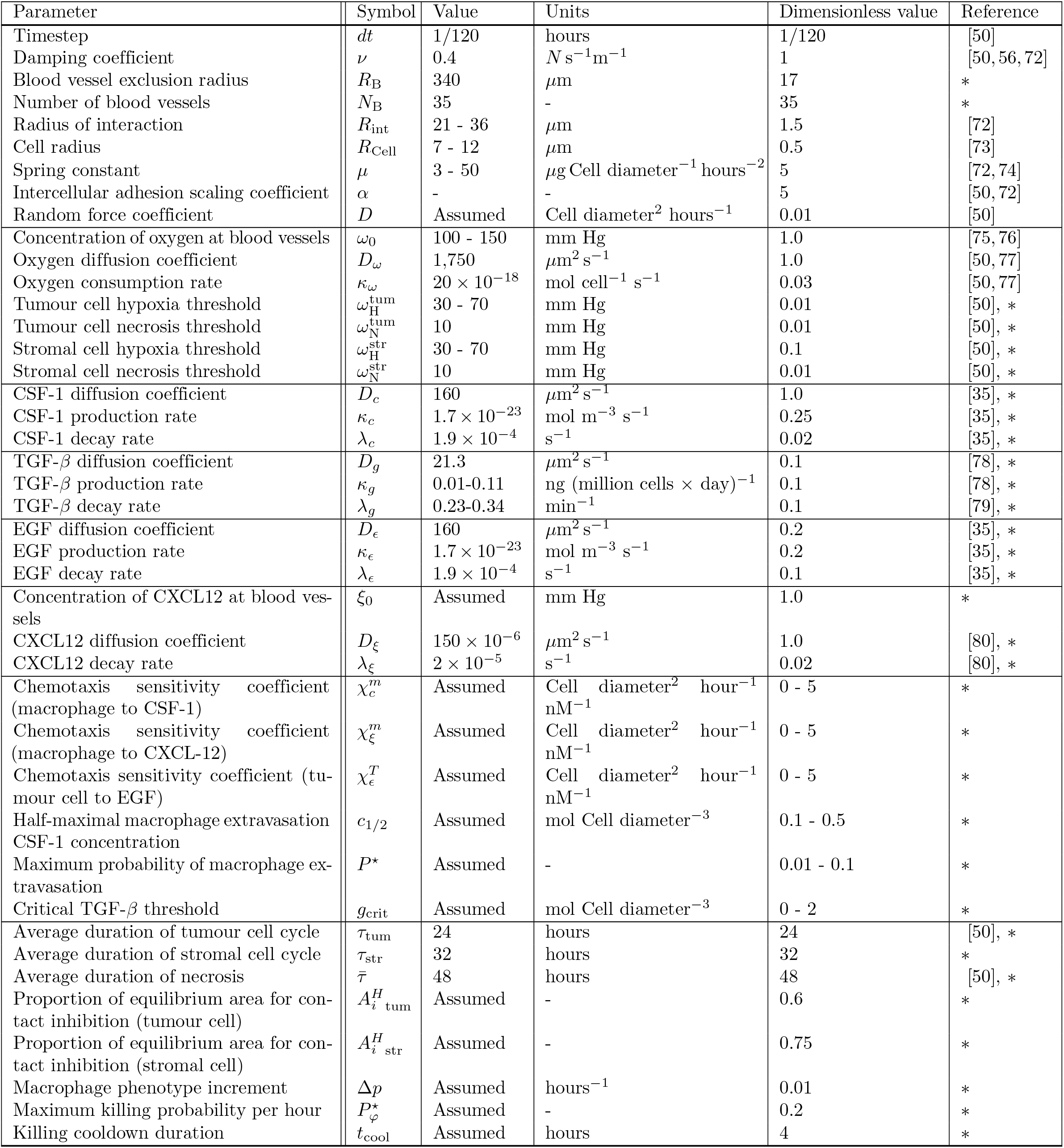
Table of parameters

Our model is implemented such that typical scales are given by:

- Length: 1 cell diameter (taken as 20*μ*m) is 1 unit of length.
- Time: 1 hour is 1 unit of time.
- Concentration: the boundary concentration of oxygen is 1.

## S2 Appendix: Tumour progression in the presence and absence of macrophages

Fig 11 shows how the presence of macrophages can alter the spatial composition and growth dynamics of a small number of tumour cells initially located at the centre of a two-dimensional square domain that contains stromal cells and a fixed number of static blood vessels. The vessels are randomly distributed in the domain but excluded from a circle of radius *R*_*B*_ centred on the initial tumour mass. Panels A and C show that, in the absence of macrophages (i.e., setting the maximum probability of extravasation *P*^⋆^ = 0), the tumour increases rapidly in size but remains as a compact mass. At much longer timescales, the tumour evolves to a steady state where the net proliferation rate of oxygen-rich cells on the tumour periphery balances the net death rate of oxygen-starved cells towards the tumour centre.

**Fig 11.**
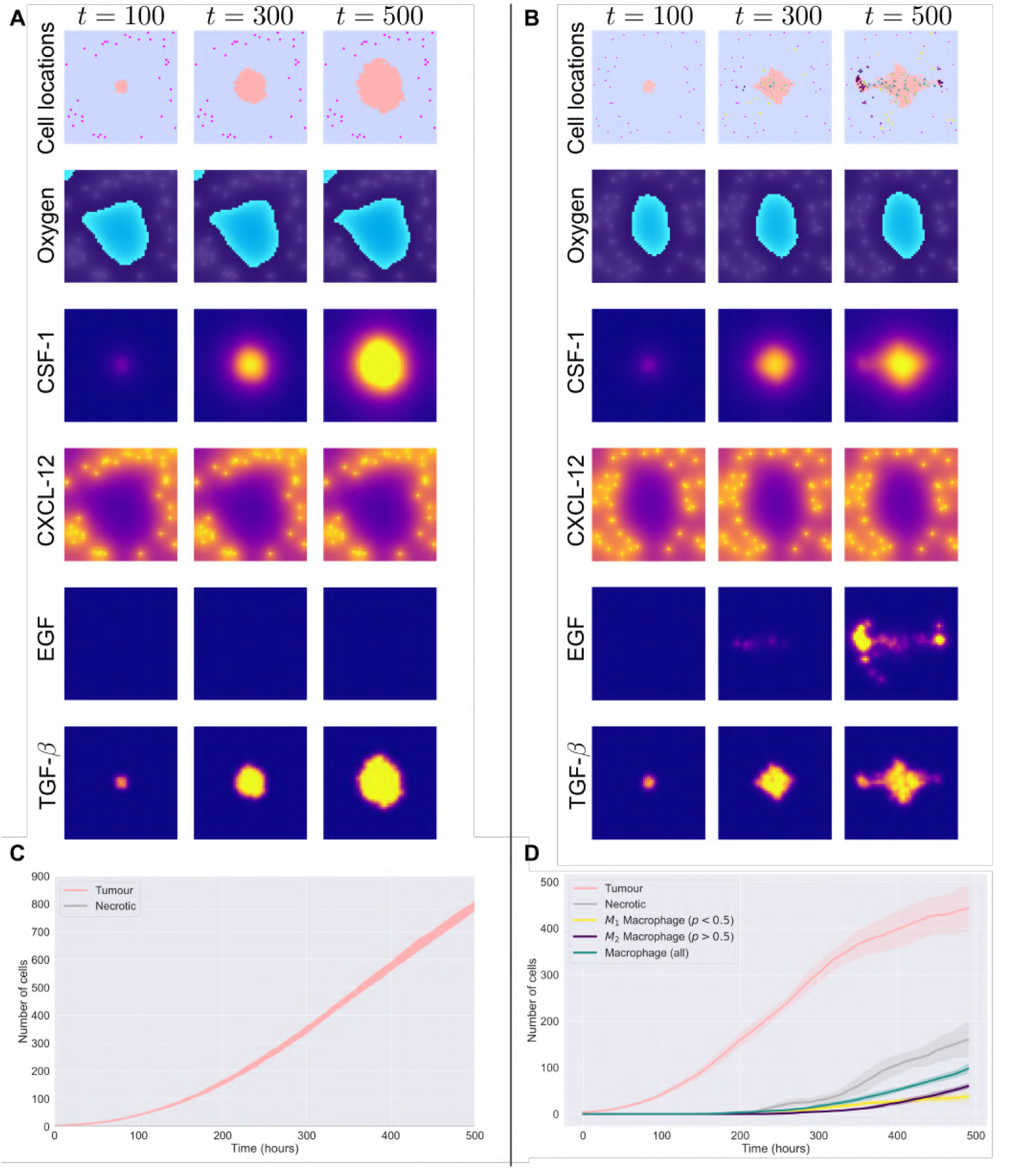
Representative output from model simulations. A, B: Spatial distributions of cells, oxygen, CSF-1, CXCL12, EGF and TGF-*β* at times *t* = 100, 300, 500 from simulations which neglect (A, *P*^⋆^ = 0) or include (B, *P*^⋆^ = 0.075) macrophage extravasation. Comparison of these plots shows how the tumour’s growth rate and spatial composition can change in the presence of macrophages. Parameter values: 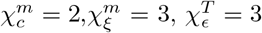, *c*_1/2_ = 0.9, *g*_*crit*_ = 0.05; all other parameters fixed at c ξ ϵ their default values (see Table 2 in S1 Appendix: Model Description). C, D: Change in numbers of tumour cells, necrotic cells, M_1_ and M_2_ macrophages, and total number of macrophages over time for the simulations presented in (A) and (B) (mean and SD from 10 realisations).

When macrophage extravasation is active (Fig 11B and Fig 11D), the ABM reproduces the qualitative behaviours outlined in Fig 2 [33]. At early times (*t* ≈ 100), CSF-1 levels are below the threshold for macrophage extravasation and the domain is devoid of macrophages. As the tumour increases in size, more CSF-1 is produced until, eventually, CSF-1 levels at the blood vessels reach the threshold for extravasation of M_1_ macrophages (*t* ≈ 200). By *t* = 300, some macrophages have infiltrated the tumour mass. Macrophages that have been exposed to sufficient levels of TGF-*β* become M_2_ and migrate back towards nearby blood vessels, in response to spatial gradients in CXCL12. The M_2_ macrophages also produce EGF which acts as a chemoattractant for the tumour cells. Thus, at *t* = 500 hours, clusters of M_2_ macrophages and tumour cells surrounding multiple blood vessels are visible in the domain.

Comparison of Fig 11A and Fig 11B, and Fig 11C and Fig 11D, reveals how the presence of macrophages can transform a tumour from a rapidly growing, compact mass to one that is slower growing and more diffuse. While the summary data presented in Fig 11D provide useful information about the tumour’s overall growth dynamics and changing cellular composition, detailed information about its morphology and spatial heterogeneity is lacking. In Fig 12 we present additional statistics generated from the spatial data at *t* = 500. Fig 12B shows the cross-PCF *g*_*TB*_(*r*) (mean and SD of 10 simulations) for the tumour cells and blood vessels in Fig 12A. There is complete exclusion between blood vessels and tumour cells up to a radius of approximately 5 cell diameters, a lengthscale characterising the minimum distance between the blood vessels and the tumour mass. By comparison, the cross-PCF *g*_*TB*_(*r*) in Fig 12D quantifies the short-range clustering of tumour cells and blood vessels in Fig 12C. The peak around *r* = 1 indicates close proximity between some blood vessels and tumour cells. The cross-PCFs provide additional information about the strong short-range clustering of macrophages with tumour cells (Fig 12E) and blood vessels (Fig 12F). Macrophages are strongly correlated with tumour cells, particularly at short lengthscales, indicating colocalisation (Fig 12E). There is also strong short-range colocalisation between macrophages and blood vessels (Fig 12F), suggesting the presence of perivascular niches containing macrophages, tumour cells and blood vessels.

**Fig 12.**
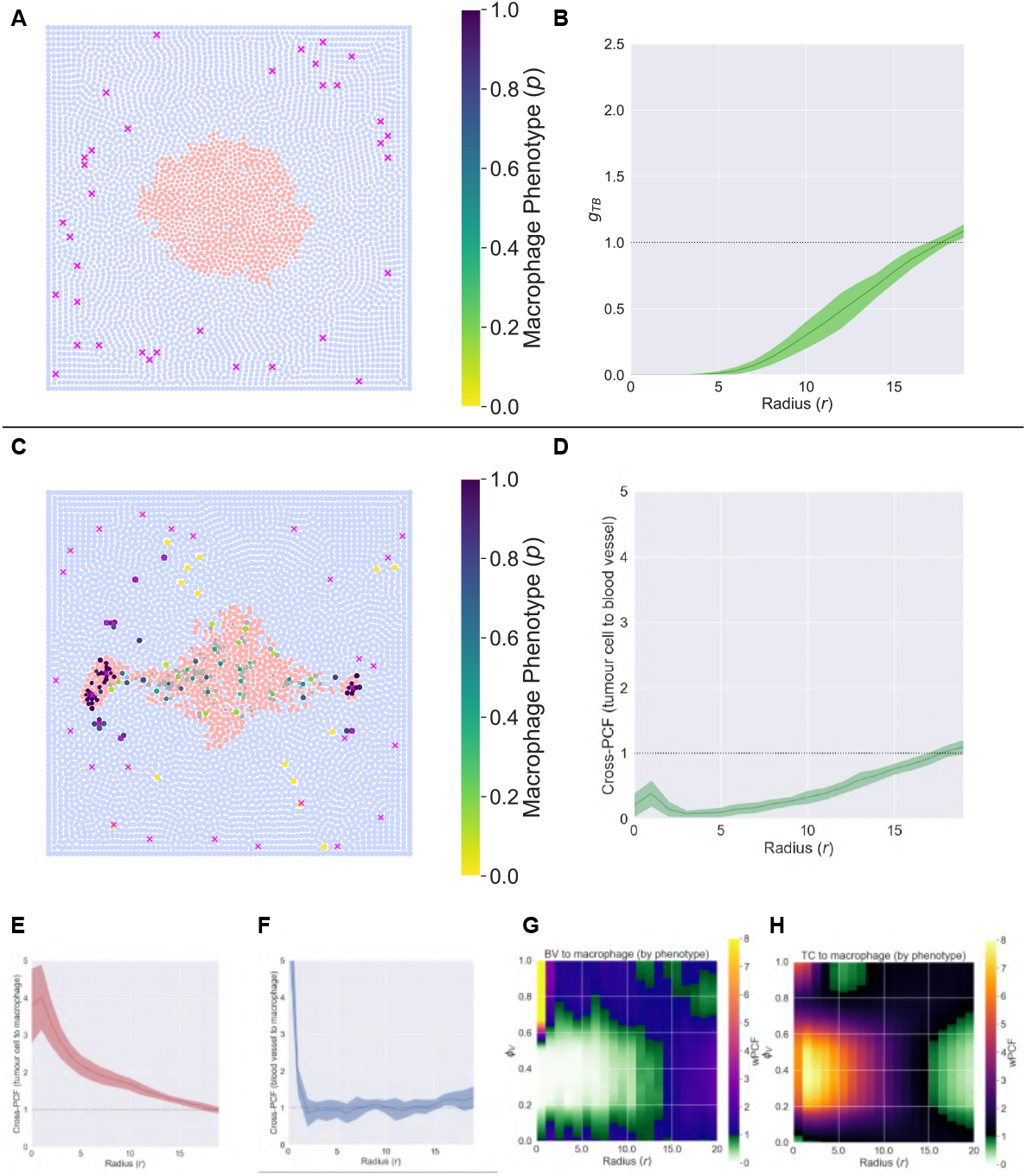
Statistical analysis of the simulation endpoints in Fig 11. A: *t* = 500 for tumour growth without macrophage extravasation. B: The cross-PCF *g*_*TB*_(*r*) for tumour cells and blood vessels in (A). No tumour cells are observed within a distance of 5 cell diameters from a blood vessel. C: *t* = 500 for tumour growth with macrophage extravasation. D: The cross-PCF *g*_*TB*_(*r*) for the tumour cells and blood vessels in (C). The cross-PCF reveals short range interactions between tumour cells and blood vessels. Comparison with (B) quantifies how the spatial distribution of tumour cells relative to the blood vessels changes in the presence of macrophages. E: The cross-PCF *g*_*TM*_ (*r*) associated with (C). The cross-PCF reveals strong short range interactions between macrophages and tumour cells. F: The cross-PCF *g*_*BM*_ (*r*) associated with (C). Short-range correlations between blood vessels and macrophages are very strong and decay rapidly with distance *r*. G: The weighted PCF *wPCF* (*r, P, B*) associated with (C). There is strong, short-range colocalisation of macrophages with *p* > 0.6 and blood vessels, while macrophages with 0.2 ⪅ *p* ⪅ 0.6 are excluded from regions of radius approximately 10-15 cell diameters surrounding blood vessels. H: The weighted PCF *wPCF* (*r, P, T*) associated with (C). Macrophages with *p* > 0.6 are strongly colocalised with tumour cells at distances 0 ≤ *r* ≤ 10, indicating their presence inside the tumour mass. Short-range colocalisation (*r* ≈ 3) is also observed for M_2_ macrophages with *p* ≥ 0.9.

The corresponding weighted PCFs, *wPCF* (*r, P, B*) and *wPCF* (*r, P, T*) (mean of 10 iterations, Fig 12G and Fig 12H) have the signature of an ‘Escape’ simulation discussed in the main text (Fig 6.

## Steady state dynamics

In this appendix we study the long term behaviour of our ABM in the absence of macrophages. We use the same (default) parameter values used to generate Fig 11, but extend the simulation time to 2000 hours.

The simulation results presented in Fig 13A show that at long times the tumour evolves to a steady state for which the rate at which oxygen-rich cells on the outer tumour boundary proliferate balances the rate at which necrotic cells in the oxygen-starved core are degraded. We have positioned the blood vessels and fixed the properties of the tumour cells (e.g., the thresholds for hypoxia and necrosis, 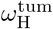 and 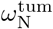, and the cell cycle duration *τ*_*i*_) so that a tumour initially located at the centre of the domain grows as a compact mass. At long times, the total number of tumour cells and necrotic cells remains approximately constant (see Figs 13B and C). Further, the tumour attains its steady state before it can spread to the surrounding blood vessels.

**Fig 13.**
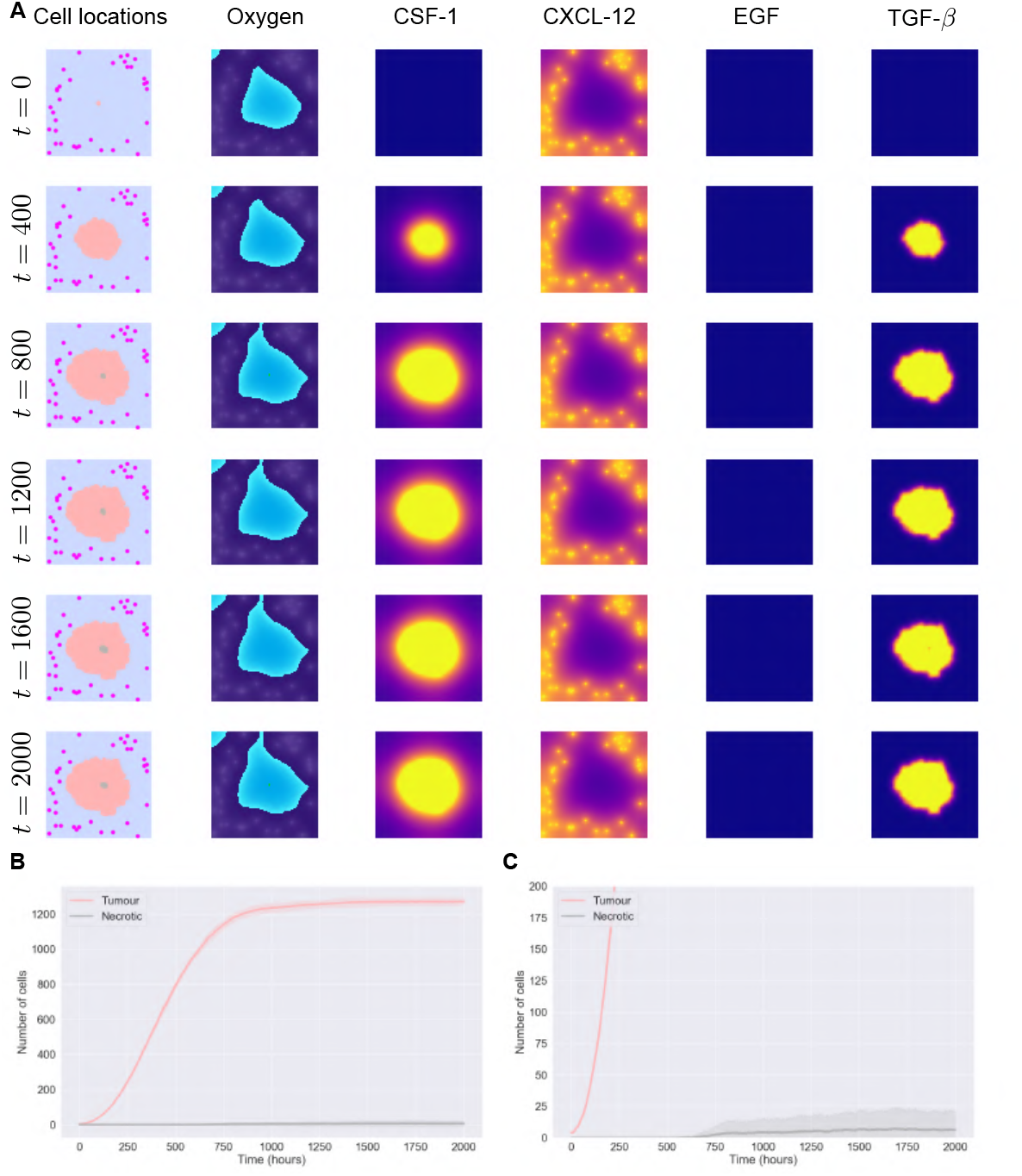
Tumour growth in the absence of macrophages. A: When no macrophages enter the simulation, the tumour grows as a compact mass in response to oxygen supplied from blood vessels. At long times, the tumour attains a steady state with a central necrotic core. At steady state, the proliferation rate of cells on the oxygen rich outer tumour boundary balances the death rate of cells in the central, oxygen-starved necrotic core. B/C: Tumour cell counts reach a steady state at approximately *t* = 1000. After approximately 750 a necrotic core forms due to hypoxia at the tumour centre. (Panel C shows a magnified view of panel B).

## S3 Appendix: Derivation of PCFs and Cross-PCFs from wPCF

In the main text, we use the wPCF to identify correlations between a continuous label (phenotype) and a categorical one (cell type). We now explain how we can extend it to study correlations between two continuous labels. Consider two labels *u* and *v* associated with each cell, which may be continuous or discrete. For two target marks *U* and *V*, the wPCF in its most general form can be written as:

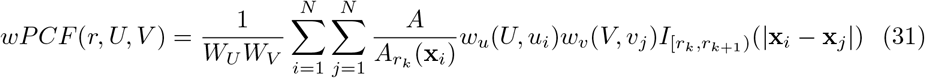

where *w*_*u*_ and *w*_*v*_ are weighting functions, and *W*_*U*_ = Σ_*i*_ *w*_*u*_(*U, u*_*i*_) and *W*_*V*_ = Σ_*i*_ *w*_*v*_(*V, v*_*i*_) are the total weights associated with each mark. The functional forms of *w*_*u*_ and *w*_*v*_ depend on several factors:

- Are *u* and *v* discrete or continuous marks?
- What are the ranges of *u* and *v*?
- At what resolution do we wish to identify correlations?

Through appropriate choices of *w*_*U*_ and *w*_*V*_, Equation (31) reduces to the ordinary PCF, the cross-PCF, or the discrete-continuous form of the wPCF that was introduced in the main text. Table 3 illustrates weighting functions which achieve these different cases.

**Table 3.** Choices of *w* which simplify the wPCF. By appropriately choosing *w*_*u*_ and *w*_*v*_, Equation (31) reduces to the wPCF presented in Equation (11), to the cross-PCF in Equation (9), or to the original definition of the PCF.

| Function name | Notation | $u$: Discrete? | $v$: Discrete? | $w_u$ | $w_v$ |
| --- | --- | --- | --- | --- | --- |
| PCF | $g(r)$ | Discrete | Discrete | $w_u(U, u_i) = \Theta(U, u_i)$ | $w_v(V, v_i) = \Theta(V, v_i)$ with $u = v$ |
| Cross-PCF | $g_{UV}(r)$ | Discrete | Discrete | $w_u(U, u_i) = \Theta(U, u_i)$ | $w_v(V, v_i) = \Theta(V, v_i)$ |
| wPCF | $wPCF(r, U, V)$ | Continuous | Discrete | $w_u(U, u_i)$ | $w_v(V, v_i) = \Theta(V, v_i)$ |
| wPCF | $wPCF(r, U, V)$ | Continuous | Continuous | $w_u(U, u_i)$ | $w_v(V, v_i)$ |

As an example, Fig 14 shows the cross-PCFs and wPCFs from Fig 5 in the main text. Through appropriate choice of the weighting function used in the wPCF, the cross-PCFs can be thought of as wPCFs in which the weighting function is piecewise constant as a function of *P*.

**Fig 14.**
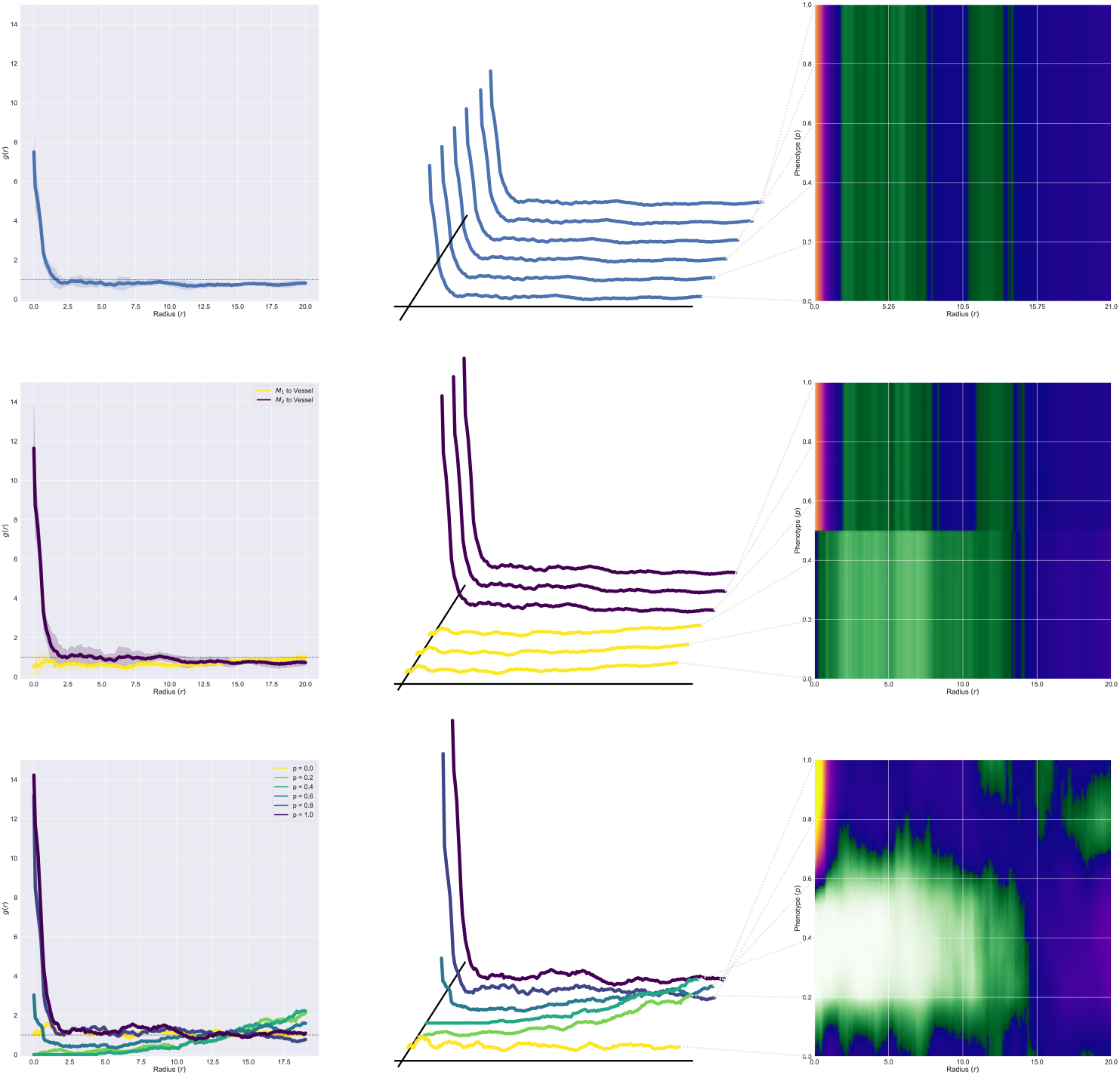
PCFs and Cross-PCFs can be seen as special cases of the wPCF. Cross- and wPCFs from Fig 5. By appropriately choosing the weighting function *w*, the wPCF returns the same results as the cross-PCF. Top row: *wPCF* (*r, P, B*) = *g*_*BM*_ (*r*), with weighting function *w*(*P, p*_*i*_) = 1 for all target phenotypes *P* and cell phenotypes *p*_*i*_. Middle row: 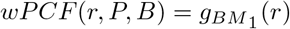 for *P* ≤ 0.5 and 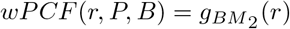 for *P* > 0.5, with weighting function *w*(*P, p*_*i*_) = 1 if *P* and *p*_*i*_ are both above/below 0.5, and 0 if they are on opposite sides. Bottom row: standard wPCF as described in Fig 5.

## S4 Appendix: Comparison of different weighting functions

In this appendix we consider different choices of the weighting, or kernel, function *w*_*p*_, and apply them to synthetic data. In Fig 15 we place 50 pink crosses uniformly along the line *y* = 1, and 1000 circles according to complete spatial randomness throughout the (2 × 2) square domain. The circles are labelled with a ‘phenotype’ *p* based on their distance from *y* = 1, such that for circle *i* the label *p*_*i*_ is given by:

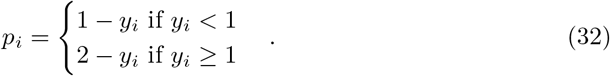

By construction, this results in two prominent values of *p* which are correlated with pink crosses at distance *r*: *p* = *r* (below *y* = 1) and *p* = 1 − *r* (above *y* = 1).

**Fig 15.**
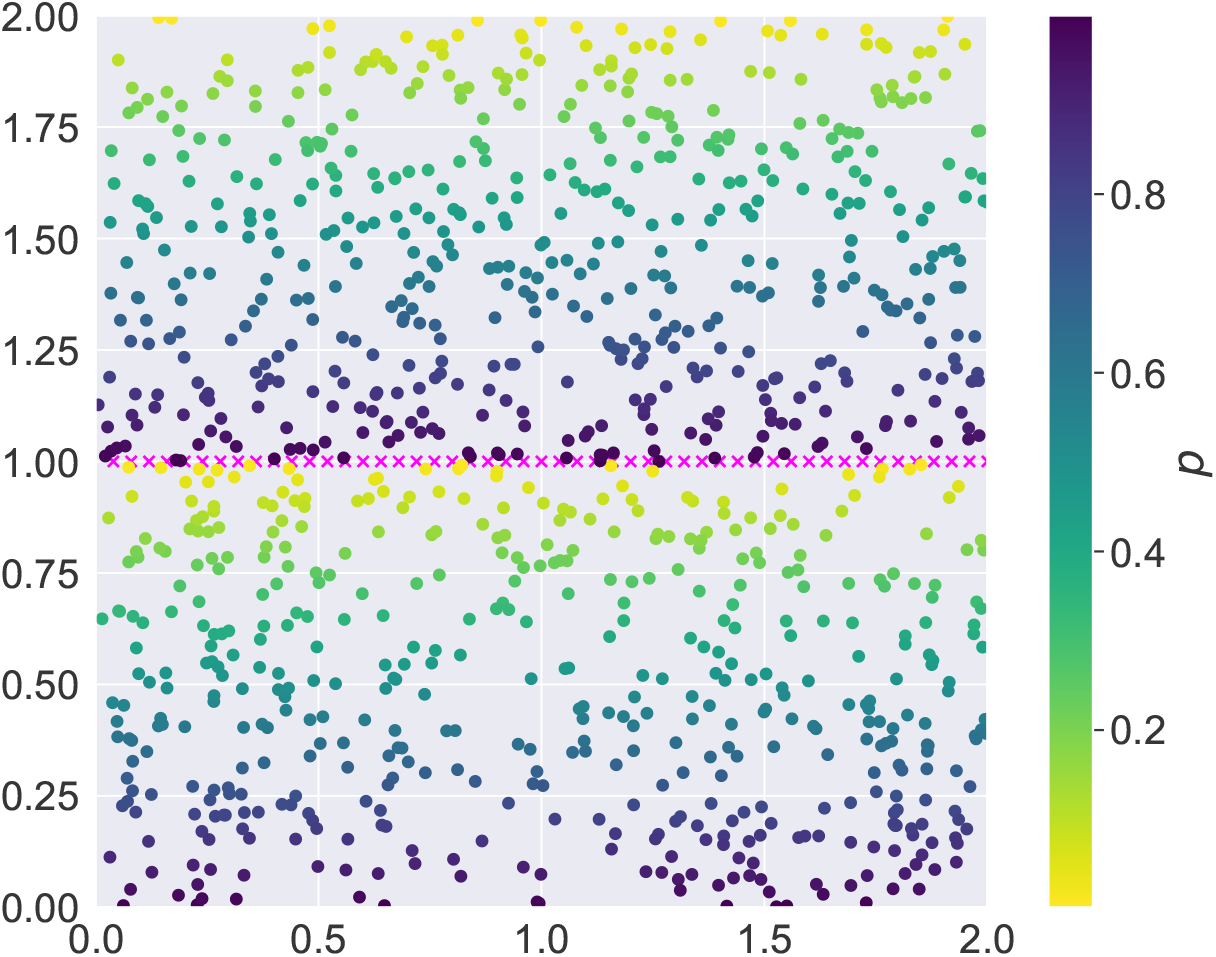
Synthetic data generated to demonstrate different weighting functions. The test data consists of 50 pink crosses (’blood vessels’ uniformly positioned along the line *y* = 1) and 1000 circles randomly positioned in the domain (‘macrophages’ with different phenotypes placed according to complete spatial randomness). Circles are labelled according to their distance from the line *y* = 1.

In Fig 16 we present wPCFs for this point cloud for different choices of the weighting function *w*_*p*_(*P, p*_*i*_) in Equation (11). In Fig 16A, we use the kernel from the main text, which has the form 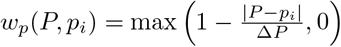, and consider different values of Δ*P*. In the main text, we fixed Δ*P* = 0.2, as it produces a kernel that is sufficiently compact to identify variations in phenotype while being broad enough to reduce noise. In Fig 16B, we fix 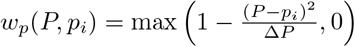, which leads to a smoother kernel and, hence, a smoother wPCF. In all cases, we present the wPCF and *w_p_*(0.5, *p_*i*_*).

**Fig 16.**
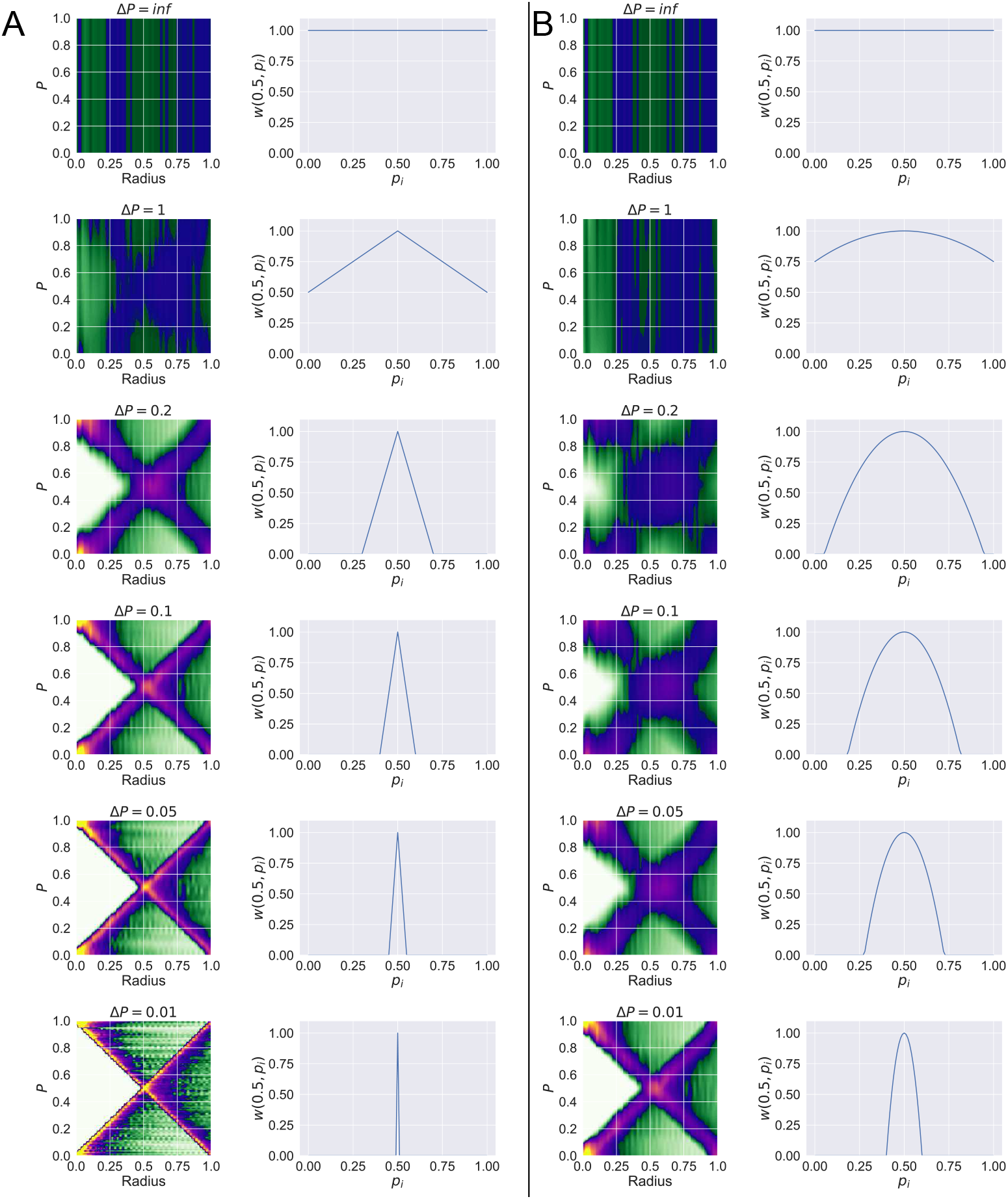
wPCFs generated from different weighting functions. *wPCF* (*r, P, B*) for the point pattern in Fig 15. Changing the shape of the weighting function adjusts the balance between signal and noise in the wPCF: the narrower the support of *w*, the more clearly the relationship between label and distance can be discerned. Using a weighting function with extremely narrow support relative to the range of labels results in more noise in the wPCF, most evident in the triangular weighting functions with Δ*P* = 0.05 and Δ*P* = 0.01. A: wPCFs generated using weighting functions of the form *w*(*P, p*) = 1 − *m|P − p|*, together with *w*(0.5, *p*) B: wPCFs generated using weighting functions of the form *w*(*P, p*) = 1 − *m*(*P − p*)^2^, together with *w*(0.5, *p*)

Fig 16 shows how the shape of the weighting function influences the balance between signal and noise in the wPCF. When the support of the weighting function is broad (e.g., Δ*P* ≥ 1), the wPCF does not identify correlations between *r* and the target phenotype *P*. On the other hand, when the support of the weighting function is narrow (e.g., Δ*P* = 0.01 for the triangular weighting function), the resulting wPCF identifies correlation well but contains a lot of noise. We conclude that the choice of weighting function can have a strong effect on the resulting wPCF, and should be chosen with care. Selecting a weighting function which is too ‘narrow’ will result in noisy wPCFs, while one that is too ‘broad’ will produce wPCFs that are unable to resolve resolution key features. In practice, the appropriate choice is likely to depend on both the distribution of labels in the data and the number of points available, in the same way that the selection of an appropriate annulus radius for the PCF must be tailored to the dataset in question.

## S5 Appendix: wPCF for comparing two continuous labels

As discussed in S3 Appendix: Derivation of PCFs and Cross-PCFs from wPCF, the wPCF can be generalised to account for relationships between points with two continuous marks (rather than a continuous mark and discrete mark). In its most general form, the PCF is given by

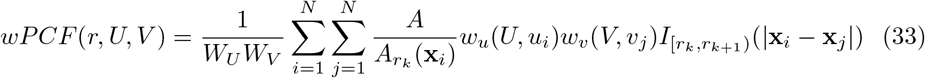

where *w*_*u*_ and *w*_*v*_ are appropriately chosen weighting functions, and *W*_*U*_ = Σ_*i*_ *w*_*u*_(*U, u*_*i*_) and *W*_*V*_ = Σ_*i*_ *w*_*v*_(*V, v*_*i*_) are the total weights associated with each mark.

## Demonstration on CSR with correlated labels

To demonstrate this, we consider the synthetic point pattern in Fig 17A. There are two different types of points: 200 circles with a continuous label *p* ∈ [0, 1], and 200 triangles with a continuous label *ψ* ∈ [0, 10], with locations distributed according to complete spatial randomness. The scales of the labels are chosen to demonstrate the ability of the wPCF to analyse marks which vary over different ranges. The location (*x*_*i*_, *y*_*i*_) of each point is chosen randomly, but their labels are defined as follows: *p*_*i*_ = *y*_*i*_ and *ψ*_*i*_ = 10(1 − *y*_*i*_).

**Fig 17.**
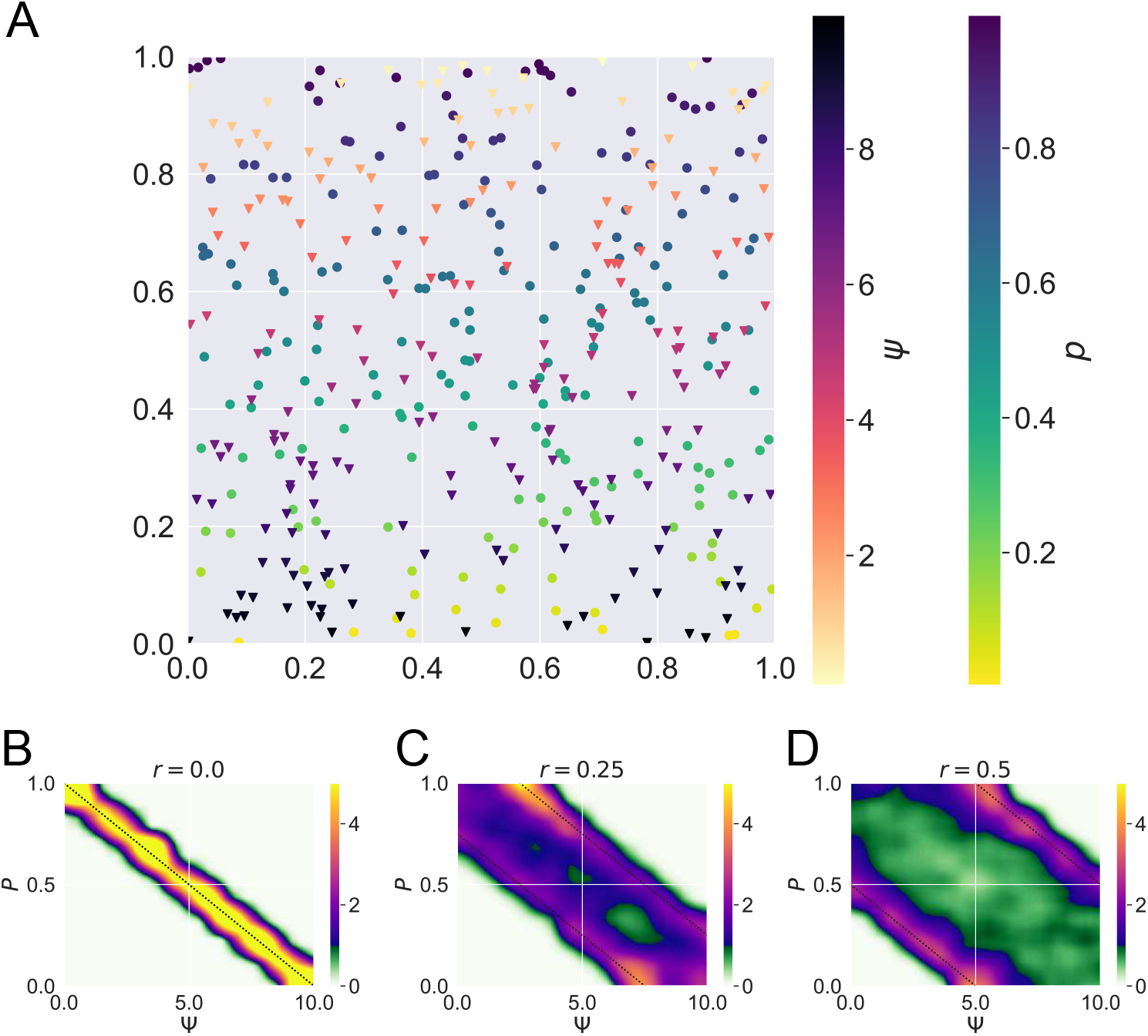
wPCF comparing two continuous labels. A: Two types of points (circles with label *p* and triangles with label *ψ*) distributed at random. Points are labelled according to their *y* coordinate, with point *i* having *p*_*i*_ = *y*_*i*_ or *ψ*_*i*_ = 10(1 − *y*_*i*_). B-D : *wPCF* (*r, P*, Ψ) for B) *r* = 0, C) *r* = 0.25, D) *r* = 0.5. Dotted black lines show the expected maximal values of the wPCF according to the construction of the point pattern.

We expect to see colocalisation of triangles with label *ψ* and circles with label *p* = 1 − 0.1*ψ*. More generally, we expect to observe strong correlation at distance *r* between triangles with a target label Ψ and circles with a target label *P* = 1 − 0.1Ψ ± *r*.

Since we must now specify two target labels, *p* = *P* and *ψ* = Ψ, and a radius, *r*, *wPCF* (*r, P*, Ψ) is a three dimensional statistic. This can be most easily visualised by considering fixed values of *r*, and observing which values of *P* and Ψ lead to higher or lower values of the wPCF. Figs 17B-D show *wPCF* (*r, P*, Ψ) for *r* = 0, 0.25, 0.5. The dotted black lines in each panel represent the lines *P* = 1 − 0.1Ψ ± *r*, and show that the wPCF successfully describes the relationships between points with label *P* and Ψ separated by distance *r*.

In this example, we take 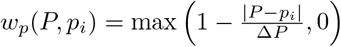 with Δ*P* = 0.1, and 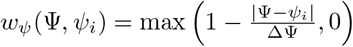 with ΔΨ = 1, to reflect the fact that the range of *ψ* is 10 times larger than that of *p*.

## Demonstration on data from Fig 6

Figs 18 and 19 show correlations between macrophages with phenotype *p*_*A*_ and those with phenotype *p*_*B*_ in the simulation endpoints shown in Fig 6 (averaged over 10 repetitions).

**Fig 18.**
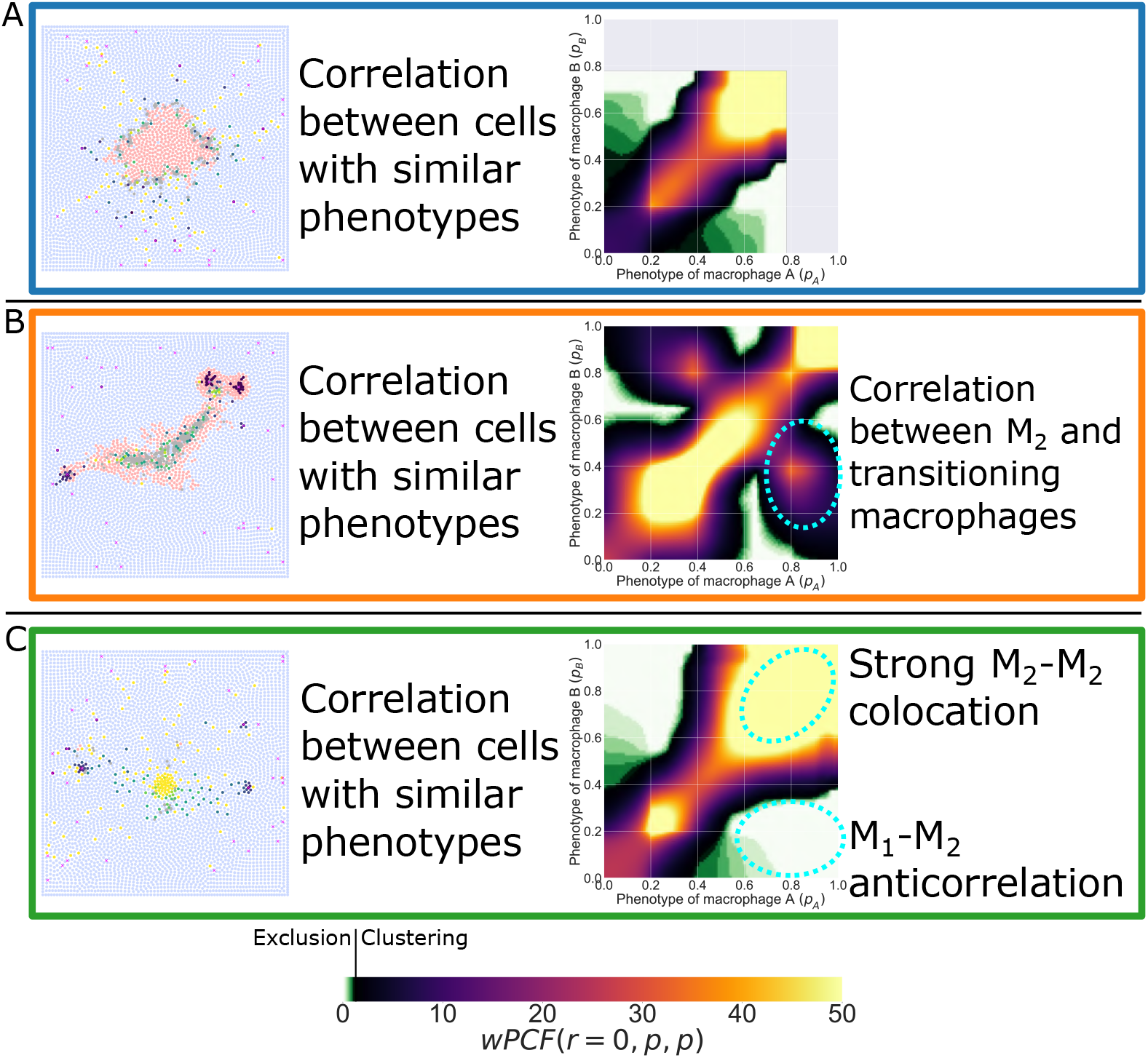
wPCF comparing two continuous labels. *wPCF* (*r* = 0, *p*_*A*_, *p*_*B*_) for simulations in Fig 6. These plots show clustering between macrophages with phenotype *p*_*A*_ and *p*_*B*_ at short distances (*r* ∈ [0, 1]).

**Fig 19.**
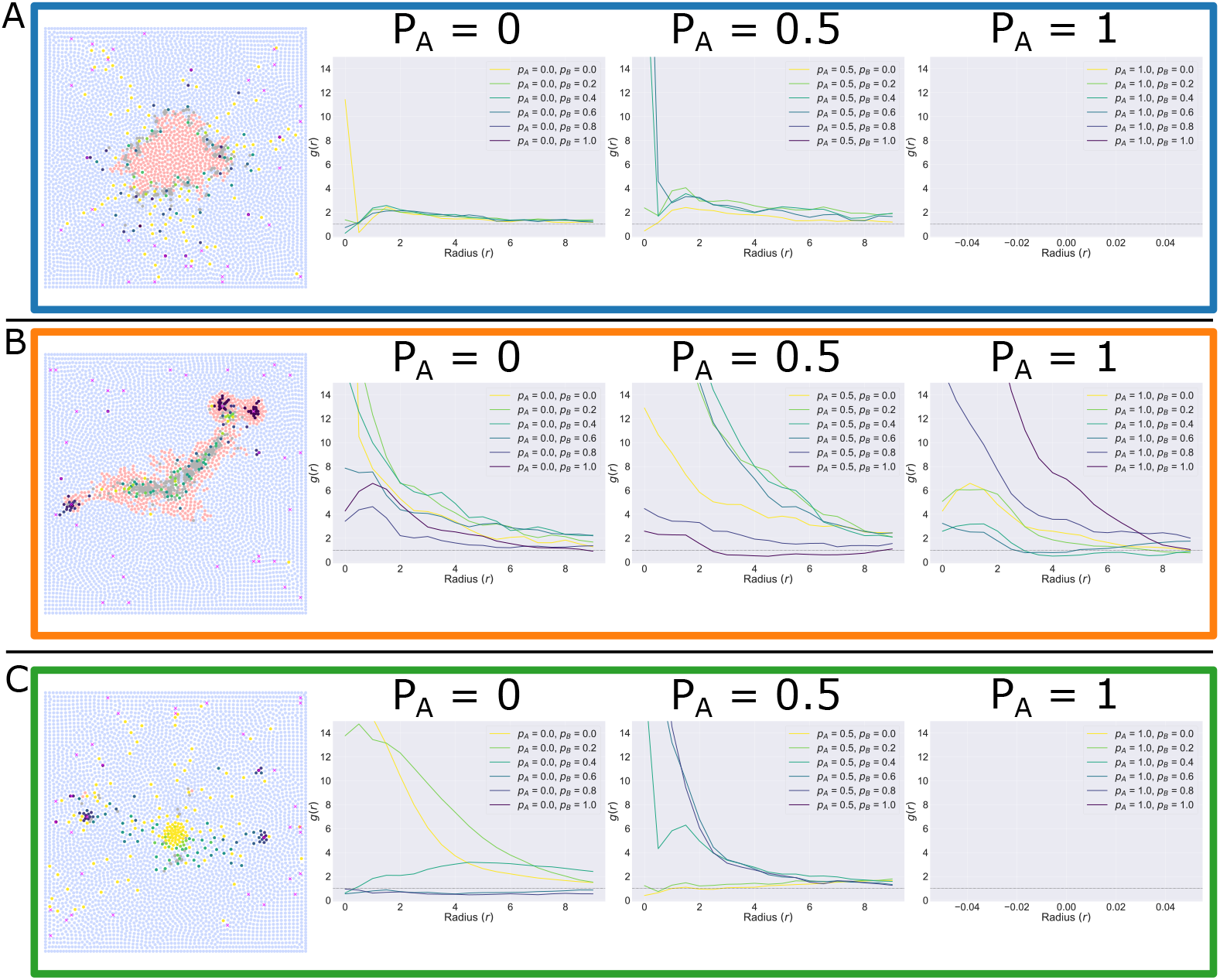
wPCF comparing two continuous labels. *wPCF* (*r, p*_*A*_, *p*_*B*_) for fixed values *p*_*A*_ ∈ [0, 0.5, 1] and *p*_*B*_ ∈ [0, 0.2, 0.4, 0.6, 0.8, 1]. As in Fig 3, visualising the wPCF for fixed values of *p*_*A*_ and *p*_*B*_ leads to lines with the familiar interpretation of a cross-PCF.

Fig 18 shows *wPCF* (*r* = 0, *p*_*A*_, *p*_*B*_). In each case, macrophages with similar phenotypes are observed within distance *r* ∈ [0, 1] much more often than spatial randomness would suggest, indicating that similar macrophages are frequently found together. We highlight two features of the simulations highlighted by *wPCF* (*r* = 0, *p*_*A*_, *p*_*B*_). Firstly, in panel B we see a peak at *p*_*A*_ = 0.8, *p*_*B*_ = 0.4 (and vice versa). This shows colocation between macrophages which are at the M_2_ end of the phenotype spectrum and macrophages which have begun to transition. Secondly, in panel C we see strong anti-correlation between M_1_ and M_2_ macrophages, emphasising that they preferentially localise in different areas of the tissue.

Fig 19 shows an alternative way of visualising *wPCF* (*r, p*_*A*_, *p*_*B*_), in which we fix specific values of *p*_*A*_ and *p*_*B*_ in order to visualise correlations at different lengthscales.

We note that the scale of the wPCF here (in some places *wPCF* > 50) suggests that these effects are likely to be strongly influenced by individual cells, since there are only small numbers of macrophages present with some combinations of *p*_*A*_ and *p*_*B*_.

## References

1. Bindea G, Mlecnik B, Tosolini M, Kirilovsky A, Waldner M, Obenauf AC, Angell H, Fredriksen T, Lafontaine L, Berger A, Bruneval P, Fridman WH, Becker C, Pagès F, Speicher MR, Trajanoski Z, Galon J. Spatiotemporal dynamics of intratumoral immune cells reveal the immune landscape in human cancer. Immunity. 2013;39(4):782–795. doi:10.1016/j.immuni.2013.10.003.

2. Singleton DC, Macann A, Wilson WR. Therapeutic targeting of the hypoxic tumour microenvironment. Nature Reviews Clinical Oncology. 2021;18(12):751–772. doi:10.1038/s41571-021-00539-4.

3. Gentles AJ, Newman AM, Liu CL, Bratman SV, Feng W, Kim D, Nair VS, Xu Y, Khuong A, Hoang CD, Diehn M, West RB, Plevritis SK, Alizadeh AA. The prognostic landscape of genes and infiltrating immune cells across human cancers. Nature Medicine. 2015;21(8):938–945. doi:10.1038/nm.3909.

4. Cortese N, Carriero R, Laghi L, Mantovani A, Marchesi F. Prognostic significance of tumor-associated macrophages: past, present and future. Seminars in Immunology. 2020;48:101408. doi:10.1016/j.smim.2020.101408.

5. Mantovani A, Marchesi F, Malesci A, Laghi L, Allaven P. Tumour-associated macrophages as treatment targets in oncology. Nature Reviews Clinical Oncology. 2017;14(7):399–416. doi:10.1038/nrclinonc.2016.217.

6. Li Z, Maeda D, Yoshida M, Umakoshi M, Nanjo H, Shiraishi K, et al. The intratumoral distribution influences the prognostic impact of CD68- and CD204-positive macrophages in non-small cell lung cancer. Lung Cancer. 2018;123:127–135. doi:10.1016/j.lungcan.2018.07.015.

7. Rakaee M, Rasmussen Busund L-T, Jamaly S, Paulsen E-E, Richardsen E, et al. Prognostic Value of Macrophage Phenotypes in Resectable Non-Small Cell Lung Cancer Assessed By Multiplex Immunohistochemistry. Neoplasia. 2019;21(3):282–293. doi:10.1016/j.neo.2019.01.005.

8. Pinto ML, Rios E, Durães C, Ribeiro R, Machado JC, Mantovani A, Barbosa MA, Carneiro F, Oliveira MJ. The Two Faces of Tumor-Associated Macrophages and Their Clinical Significance in Colorectal Cancer. Frontiers in Immunology. 2019;10:1–12. doi:10.3389/fimmu.2019.01875.

9. López-Janeiro Á, Padilla-Ansala C, de Andrea CE, Hardisson D, Melero I. Prognostic value of macrophage polarization markers in epithelial neoplasms and melanoma. A systematic review and meta-analysis. Modern Pathology. 2020;33(8):1458–1465. doi:10.1038/s41379-020-0534-z.

10. Jackute J, Zemaitis M, Pranys D, Sitkauskiene B, Miliauskas S, Vaitkiene S, Sakalauskas R. Distribution of M1 and M2 macrophages in tumor islets and stroma in relation to prognosis of non-small cell lung cancer. BMC Immunology. 2018;19(3):1–13. doi:10.1186/s12865-018-0241-4.

11. Murray PJ, Allen JE, Biswas SK, Fisher EA, Gilroy DW, Goerdt S, Gordon S, Hamilton JA, Ivashkiv LB, Lawrence T, Locati M, Mantovani A, Martinez FO, Mege J-L, Mosser DM, Natoli G, Saeij JP, Schultze JL, Shirey KA, Sica A, Suttles J, Udalova I, van Ginderachter JA, Vogel SN, Wynn TA. Macrophage Activation and Polarization: Nomenclature and Experimental Guidelines. Immunity. 2014;41(1):14–20. doi:10.1016/j.immuni.2014.06.008.

12. Huang YK, Wang M, Sun Y, Di Costanzo N, Mitchell C, Achuthan A, Hamilton JA, Busuttil RA, Boussioutas A. Macrophage spatial heterogeneity in gastric cancer defined by multiplex immunohistochemistry. Nature Communications. 2019;10(1):1–15. doi:10.1038/s41467-019-11788-4.

13. Giesen C, Wang HAO, Schapiro D, Zivanovic N, Jacobs A, Hattendorf B, Schüffler PJ, Golimund D, Buhmann JM, Brandt S, Varga Z, Wild PJ, Günther D, Bodenmiller B Highly multiplexed imaging of tumor tissues with subcellular resolution by mass cytometry. Nature Methods. 2014;11(4):417–422. doi:10.1038/nmeth.2869.

14. Kuett L, Catena R, Özcan A, Plüss A, Cancer Grand Challenges IMAXT Consortium, Schraml P, Moch H, de Souza N, Bodenmiller B. Three-dimensional imaging mass cytometry for highly multiplexed molecular and cellular mapping of tissues and the tumor microenvironment. Nature Cancer. 2022;3(1):122–133. doi:10.1038/s43018-021-00301-w.

15. Wilsom CM, Ospina OE, Townsend MK, Nguyen J, Moran Segura C, Schildkraut JM, Tworoger SS, Peres LC, Fridley BL. Challenges and Opportunities in the Statistical Analysis of Multiplex Immunofluorescence Data. Cancers. 2021;13,3031. doi:10.3390/cancers13123031.

16. Ben-Said M. Spatial point-pattern analysis as a powerful tool in identifying pattern-process relationships in plant ecology: an updated review. Ecological Processes. 2021;10(56). doi:10.1186/s13717-021-00314-4.

17. Szmyt J. Spatial statistics in ecological analysis: from indices to functions. Silva Fennica. 2014;48(1). doi:10.14214/sf.1008.

18. Wälder O, Stoyan D. On Variograms in Point Process Statistics. Biometrical Journal. 1996;38(8):895–905. doi:10.1002/bimj.4710380802.

19. Beisbart C, Kerscher M, Mecke K. Mark Correlations: Relating Physical Properties to Spatial Distributions. In: Mecke K, Stoyan D, editors. Morphology of Condensed Matter. Springer, Berlin, Heidelberg; 2002. pp. 358–390. doi:10.1007/3-540-45782-8_15.

20. Beisbart C, Kerscher M. Luminosity- and morphology-depdendent clustering of galaxies. The Astrophysical Journal. 2000;545:6–25. doi:10.1086/317788.

21. Stoyan D, Stoyan H. Fractals, Random Shapes and Point Fields: Methods of Geometrical Statistics. Chichester, Wiley; 1994.

22. Gavrikov V, Stoyan D. The use of marked point processes in ecological and environmental forest studies. Environmental and Ecological Statistics. 1995;2(4):331–344. doi:10.1007/BF00569362.

23. Stoyan D, Wälder O. On Variograms in Point Process Statistics, II: Models of Markings and Ecological Interpretation. Biometrical Journal. 2000;42(2):171–187. doi:10.1002/(SICI)1521-4036(200005)42:2<171::AID-BIMJ171>3.0.CO;2-L.

24. Klowss JJ, Browning AP, Murphy RJ, Carr EJ, Plank MJ, Gunasingh G, Haass NK, Simpson MJ. A stochastic mathematical model of 4D tumour spheroids with real-time fluorescent cell cycle labelling. Journal of the Royal Society Interface. 2022;19:20210903. doi:10.1098/rsif.2021.0903.

25. Binder BJ, Simpson MJ. Quantifying spatial structure in experimental observations and agent-based simulations using pair-correlation functions. Physical Review E - Statistical, Nonlinear, and Soft Matter Physics. 2013;88(2):1–10. doi:10.1103/PhysRevE.88.022705.

26. Agnew DJG, Green JEF, Brown TM, Simpson MJ, Binder BJ. Distinguishing between mechanisms of cell aggregation using pair-correlation functions. Journal of Theoretical Biology. 2014;352:16–23. doi:10.1016/j.jtbi.2014.02.033.

27. Gavagnin E, Owen JP, Yates CA. Pair correlation functions for identifying spatial correlation in discrete domains. Physical Review E. 2018;97(6). doi:10.1103/PhysRevE.97.062104.

28. Johnston ST, Crampin EJ. Corrected pair correlation functions for environments with obstacles. Physical Review E. 2019;99(3):1–19. doi:10.1103/PhysRevE.99.032124.

29. Browning AP, McCue SW, Binny RN, Plank MJ, Shah ET, Simpson MJ. Inferring parameters for a lattice-free model of cell migration and proliferation using experimental data. Journal of Theoretical Biology. 2018;437:251–260. doi:10.1016/j.jtbi.2017.10.032.

30. Vipond O, Bull JA, Macklin PS, Tillmann U, Pugh CW, Byrne HM, Harrington HA. Multiparameter persistent homology landscapes identify immune cell spatial patterns in tumors. Proceedings of the National Academy of Sciences. 2021;118(41):e2102166118. doi:10.1073/pnas.2102166118.

31. Fozard JA, Kirkham GR, Buttery LDK, King JR, Jensen OE, Byrne HM. Techniques for analysing pattern formation in populations of stem cells and their progeny. BMC Bioinformatics. 2011;12:396. doi:10.1186/1471-2105-12-396.

32. Dini S, Binder BJ, Green JEF. Understanding interactions between populations: Individual based modelling and quantification using pair correlation functions. Journal of Theoretical Biology. 2018;439:50–64. doi:S0022519317305210.

33. Arwert EN, Harney AS, Entenberg D, Wang Y, Sahai E, Pollard JW, et al. A Unidirectional Transition from Migratory to Perivascular Macrophage Is Required for Tumor Cell Intravasation. Cell Reports. 2018;23(5):1239–1248. doi:10.1016/j.celrep.2018.04.007.

34. Harney AS, Arwert EN, Entenberg D, Wang Y, Guo P, Qian BZ, et al. Real-Time Imaging Reveals Local, Transient Vascular Permeability, and Tumor Cell Intravasation Stimulated by TIE2hi Macrophage-Derived VEGFA. Cancer Discovery. 2015;5(9):932–943. doi:10.1158/2159-8290.CD-15-0012.

35. Elitas M, Zeinali S. Modeling and Simulation of EGF-CSF-1 pathway to Investigate Glioma - Macrophage Interaction in Brain Tumors. International Journal of Cancer Studies & Research (IJCR). 2016; p. 1–8.

36. Knútsdóttir H, Pálsson E, Edelstein-Keshet L. Mathematical model of macrophage-facilitated breast cancer cells invasion. Journal of Theoretical Biology. 2014;357:184–199. doi:10.1016/j.jtbi.2014.04.031.

37. Knútsdóttir H, Condeelis JS, Pálsson E. 3-D individual cell based computational modeling of tumor cell–macrophage paracrine signaling mediated by EGF and CSF-1 gradients. Integrative Biology. 2016;8(1):104–119. doi:10.1039/C5IB00201J.

38. Owen MR, Sherratt JA. Mathematical modelling of macrophage dynamics in tumours. Mathematical Models and Methods in Applied Sciences. 1999;9(4):513–539. doi:10.1142/S0218202599000270.

39. Kelly CE, Leek RD, Byrne HM, Cox SM, Harris AL, Lewis CE. Modelling Macrophage Infiltration into Avascular Tumours. Journal of Theoretical Medicine. 2002;4(1):21–38. doi:10.1080/10273660290015242.

40. Suveges S, Eftimie R, Trucu D. Directionality of Macrophages Movement in Tumour Invasion: A Multiscale Moving-Boundary Approach. Bulletin of Mathematical Biology. 2020;82(12):1–50. doi:10.1007/s11538-020-00819-7.

41. Li X, Jolly MK, George JT, Pienta KJ, Levine H. Computational modeling of the crosstalk between macrophage polarization and tumor cell plasticity in the tumor microenvironment. Frontiers in Oncology. 2019;9(JAN):1–12. doi:10.3389/fonc.2019.00010.

42. Mahlbacher G, Curtis LT, Lowengrub J, Frieboes HB. Mathematical modeling of tumor-associated macrophage interactions with the cancer microenvironment. Journal for ImmunoTherapy of Cancer. 2018;6(1):1–17. doi:10.1186/s40425-017-0313-7.

43. Webb SD, Owen MR, Byrne HM, Murdoch C, Lewis CE. Macrophage-based anti-cancer therapy: Modelling different modes of tumour targeting. Bulletin of Mathematical Biology. 2007;69(5):1747–1776. doi:10.1007/s11538-006-9189-2.

44. Cess CG, Finley SD. Multi-scale modeling of macrophage—T cell interactions within the tumor microenvironment. vol. 16; 2020. Available from: http://dx.doi.org/10.1371/journal.pcbi.1008519.

45. Curtis LT, Sebens S, Frieboes HB. Modeling of tumor response to macrophage and T lymphocyte interactions in the liver metastatic microenvironment. Cancer Immunology, Immunotherapy. 2021;70(5):1475–1488. doi:10.1007/s00262-020-02785-4.

46. den Breems NY, Eftimie R. The re-polarisation of M2 and M1 macrophages and its role on cancer outcomes. Journal of Theoretical Biology. 2016;390:23–39. doi:10.1016/j.jtbi.2015.10.034.

47. Eftimie R. Investigation into the role of macrophages heterogeneity on solid tumour aggregations. Mathematical Biosciences. 2020;322(March 2019):108325. doi:10.1016/j.mbs.2020.108325.

48. El-Kenawi A, Gatenbee C, Robertson-Tessi M, Bravo R, Dhillon J, et al. Acidity promotes tumour progression by altering macrophage phenotype in prostate cancer. British Journal of Cancer. 2019;121(7):556–566. doi:10.1038/s41416-019-0542-2.

49. Dunn GP, Old LJ, Schreiber RD. The three Es of cancer immunoediting. Annu. Rev. Immunol. 2004;22:329–60. doi:10.1146/annurev.immunol.22.012703.104803.

50. Bull JA, Mech F, Quaiser T, Waters SL, Byrne HM. Mathematical modelling reveals cellular dynamics within tumour spheroids. PLOS Computational Biology. 2020;16(8):e1007961. doi:10.1371/journal.pcbi.1007961.

51. Pitt-Francis J, Pathmanathan P, Bernabeu MO, Bordas R, Cooper J, Fletcher AG, et al. Chaste: A test-driven approach to software development for biological modelling. Computer Physics Communications. 2009;180(12):2452–2471. doi:10.1016/j.cpc.2009.07.019.

52. Mirams GR, Arthurs CJ, Bernabeu MO, Bordas R, Cooper J, Corrias A, et al. Chaste: An Open Source C++ Library for Computational Physiology and Biology. PLoS Computational Biology. 2013;9(3). doi:10.1371/journal.pcbi.1002970.

53. Cooper F, Baker R, Bernabeu M, Bordas R, Bowler L, Bueno-Orovio A, et al. Chaste: Cancer, Heart and Soft Tissue Environment. Journal of Open Source Software. 2020;5(47):1848. doi:10.21105/joss.01848.

54. Scott JG, Fletcher AG, Anderson ARA, Maini PK. Spatial Metrics of Tumour Vascular Organisation Predict Radiation Efficacy in a Computational Model. PLoS Computational Biology. 2016;12(1). doi:10.1371/journal.pcbi.1004712.

55. Greenspan HP. Models for the Growth of a Solid Tumor by Diffusion. Studies in Applied Mathematics. 1972;51(4):317–340. doi:10.1002/sapm1972514317.

56. Pathmanathan P, Cooper J, Fletcher A, Mirams G, Murray P, Osborne JM, et al. A computational study of discrete mechanical tissue models. Physical biology. 2009;6(3):036001. doi:10.1088/1478-3975/6/3/036001.

57. Colling R, Pitman H, Oien K, Rajpoot N, Macklin P, CM-Path AI in Histopathology Working Group, Snead D, Sackville T, Verrill C. Artificial intelligence in digital pathology: a roadmap to routine use in clinical practice. Journal of Pathology. 2019;249:143–150. doi:10.1002/path.5310.

58. Hakkoum H, Abnane I, Idri A. Interpretability in the medical field: A systematic mapping and review study. Applied Soft Computing. 2022;117:108391. doi:10.1016/j.asoc.2021.108391.

59. Stolz BJ, Kaeppler J, Markelc B, Mech F, Lipsmeier F, Muschel RJ, Byrne HM, Harrington HA. Multiscale Topology Characterises Dynamic Tumour Vascular Networks. arXiv preprint. 2020:2008.08667 doi:10.48550/arXiv.2008.08667.

60. Bull JA, Macklin PS, Quaiser T, Braun F, Waters SL, Pugh CW, Byrne HM. Combining multiple spatial statistics enhances the description of immune cell localisation within tumours. Scientific Reports. 2020;10:18624. doi:10.1038/s41598-020-75180-9.

61. Failmezger H, Muralidhar S, Rullan A, de Andrea CE, Sahai E, Yuan Y. Topological Tumor Graphs: A Graph-Based Spatial Model to Infer Stromal Recruitment for Immunosuppression in Melanoma Histology. Cancer Research. 2020;256(4):1199–1209. doi:10.1158/0008-5472.CAN-19-2268.

62. Galon J, Mlecnik B, Bindea G, Andell HK, Berger A, et al. Towards the introduction of the ‘Immunoscore’ in the classification of malignant tumours. Journal of Pathology. 2014;232(2):199–209. doi:10.1002/path.4287.

63. AbdulJabbar K, Ahmed Raza SE, Rosenthal R, Jamal-Hanjani M, Veeriah S, et al. Geospatial immune variability illuminates differential evolution of lung adenocarcinoma. Nature Medicine. 2020;26(7):1054–1062. doi:10.1038/s41591-020-0900-x.

64. Owen MR, Byrne HM, Lewis CE. Mathematical modelling of the use of macrophages as vehicles for drug delivery to hypoxic tumour sites. Journal of Theoretical Biology. 2004;226(4):377–391. doi:10.1016/j.jtbi.2003.09.004.

65. Rockne RC, Hawkins-Daruud A, Swanson KR, Sluka JP, Glazier JA, Macklin P, Hormuth II DA, Jarrett AM, Lima EABF, Oden JT, Biros G, Yankeelov TE, Curtius K, Al Bakir I, Wodarz D, Komarova N, Aparicio L, Bordyuh M, Rabadan R, Finley SD, Enderling H, Caudell J, Moros EG, Anderson ARA, Gatenby RA, Kaznatcheev A, Jeavons P, Krishnan N, Pelesko J, Wadhwa RR, Yoon N, Nichol D, Marusyk A, Hinczewski M, Scott JG. The 2019 mathematical oncology roadmap. Physical Biology. 2019;16:041005. doi:10.1088/1478-3975/ab1a09.

66. Bull JA, Byrne HM. The Hallmarks of Mathematical Oncology. Proceedings of the IEEE. 2022. In press. doi:10.1109/JPROC.2021.3136715.

67. Norton KA, Jin K, Popel AS. Modeling triple-negative breast cancer heterogeneity: Effects of stromal macrophages, fibroblasts and tumor vasculature. Journal of Theoretical Biology. 2018;452:56–68. doi:10.1016/j.jtbi.2018.05.003.

68. Wyckoff J, Wang W, Lin EY, Wang Y, Pixley F, Stanley ER, et al. A paracrine loop between tumor cells and macrophages is required for tumor cell migration in mammary tumors. Cancer Research. 2004; p. 7022–7029. doi:10.1158/0008-5472.CAN-04-1449.

69. Wyckoff JB, Wang Y, Lin EY, Li JF, Goswami S, Stanley ER, et al. Direct visualization of macrophage-assisted tumor cell intravasation in mammary tumors. Cancer Research. 2007;67(6):2649–2656. doi:10.1158/0008-5472.CAN-06-1823.

70. Drasdo D, Höhme S. A single-cell-based model of tumor growth in vitro: monolayers and spheroids. Physical Biology. 2005;2(3):133–147. doi:10.1088/1478-3975/2/3/001.

71. Meineke FA, Potten CS, Loeffler M. Cell migration and organization in the intestinal crypt using a lattice-free model. Cell Proliferation. 2001;34(4):253–266. doi:10.1046/j.0960-7722.2001.00216.x.

72. Osborne JM, Fletcher AG, Pitt-Francis JM, Maini PK, Gavaghan DJ. Comparing individual-based approaches to modelling the self-organization of multicellular tissues. PLoS Computational Biology. 2017;13(2):1–34. doi:10.1371/journal.pcbi.1005387.

73. Laget S, Broncy L, Hormigos K, Dhingra DM, BenMohamed F, Capiod T, et al. Technical insights into highly sensitive isolation and molecular characterization of fixed and live circulating tumor cells for early detection of tumor invasion. vol. 12; 2017.

74. Dunn SJ, Näthke IS, Osborne JM. Computational models reveal a passive mechanism for cell migration in the crypt. PLoS ONE. 2013;8(11):1–18. doi:10.1371/journal.pone.0080516.

75. Mueller-Klieser WF, Sutherland RM. Oxygen tensions in multicell spheroids of two cell lines. British Journal of Cancer. 1982;45(2):256–264. doi:10.1038/bjc.1982.41.

76. Grimes DR, Kelly C, Bloch K, Partridge M. A method for estimating the oxygen consumption rate in multicellular tumour spheroids. Journal of the Royal Society Interface. 2014;11(92). doi:10.1098/rsif.2013.1124.

77. Schaller G, Meyer-Hermann M. Multicellular tumor spheroid in an off-lattice Voronoi-Delaunay cell model. Physical Review E - Statistical, Nonlinear, and Soft Matter Physics. 2005;71(5):1–16. doi:10.1103/PhysRevE.71.051910.

78. Albro MB, Nims RJ, Cigan AD, Yeroushalmi KJ, Alliston T, Hung CT, Ateshian GA. Accumulation of Exogenous Activated TGF-*β* in the Superficial Zone of Articular Cartilage. Biophysical Journal. 2013;104(8):1794–1804. doi:10.1016/j.bpj.2013.02.052.

79. Wakefield LM, Winokur TS, Hollands RS, Christopherson K, Levinson AD, Sporn MB. Recombinant Latent Transforming Growth Factor *β*1 Has a Longer Plasma Half-Life in Rats than Active Transforming Growth Factor *β*1, and a Different Tissue Distribution. The Journal of Clinical Investigation. 1990;86(6):1976–1984. doi:10.1172/JCI114932.

80. Chang SL, Cavnar SP, Luker KE, Takayama S, Luker GD, Linderman JJ. Cell, Isoform, and Environment Factors Shape Gradients and Modulate Chemotaxis. PLoS ONE. 2015;10(4):e0123450. doi:10.1371/journal.pone.0123450.

81. Ghaffarizadeh A, Heiland R, Friedman SH, Shannon MM, Macklin P. PhysiCell: An open source physics-based cell simulator for 3-D multicellular systems. PLoS Computational Biology. 2018;14(2):1–34. doi:10.1371/journal.pcbi.1005991.

82. Bravo RR, Baratchart E, West J, Schenck RO, Miller AK, Gallaher J, Gatenbee CD, Basanta D, Robertson-Tessi R, Anderson ARA. Hybrid Automata Library: A flexible platform for hybrid modeling with real-time visualization. PLoS Computational Biology. 2020;16(3):1–28. doi:10.1371/journal.pcbi.1007635.

83. Leschiera E, Lorenzi T, Shen S, Almeida L, Audebert C. A mathematical model to study the impact of intra-tumour heterogeneity on anti-tumour CD8+ T cell immune response. Journal of Theoretical Biology. 2022;538:111028. doi:10.1016/j.jtbi.2022.111028.

84. Swat MH, Thomas GL, Belmonte JM, Shirinifard A, Hmeljak D, Glazier JA. Multi-Scale Modeling of Tissues Using CompuCell3D. Methods in Cell Biology. 2012;110:325–366. doi:10.1016/B978-0-12-388403-9.00013-8.

